# Systematic analysis of microtubule plus-end networks defines EB-cargo complexes critical for mitosis in budding yeast

**DOI:** 10.1101/2022.09.08.507099

**Authors:** Nikolay Kornakov, Stefan Westermann

**Affiliations:** Department of Molecular Genetics I, Faculty of Biology, Center of Medical Biotechnology, University of Duisburg-Essen, Universitätsstrasse 2, 45141 Essen, Germany

**Keywords:** Microtubule bundles, Spindle organization, Microtubule crosslinkers, Kinetochore organization, End-Binding Proteins

## Abstract

Microtubules are ubiquitous cytoskeletal polymers with essential functions in chromosome segregation, intracellular transport and cellular morphogenesis. End-binding proteins (EBs) form the nodes of intricate microtubule plus-end interaction networks. Which EB binding partners are most critical for cell division, and how cells manage to organize a microtubule cytoskeleton in the absence of an EB protein, are open questions. Here we demonstrate that the budding yeast EB protein Bim1 executes its key mitotic functions as part of two cargo complexes-Bim1-Kar9 in the cytoplasm and Bim1-Cik1-Kar3 in the nucleus. Lack of Bim1-Kar9 during spindle orientation is compensated by accumulation of the CLIP-170 homolog Bik1 on the lattice of long cytoplasmic microtubules, which upregulates the Dynein-Dynactin nuclear migration pathway. In the nucleus a Bim1-Bik1-Cik1-Kar3 complex acts during initial metaphase spindle assembly and supports sister chromatid bi-orientation. Lack of Bim1 alters spindle association timing and the level of the microtubule crosslinkers Ase1/PRC1 and Slk19, which become essential for bi-orientation. Engineered plus-end targeting of Kinesin-14 Cik1-Kar3 efficiently restores major spindle-related *bim1τι* phenotypes. In addition to defining the key Bim1-cargo complexes our study also reveals compensatory mechanisms that allow cells to proliferate in the absence of Bim1.

## Introduction

Microtubules are polar cytoskeletal filaments assembled from heterodimers of α- and β-tubulin. Their dynamics are described by a non-equilibrium behavior in which individual filaments stochastically switch between phases of polymerization and depolymerization. This phenomenon, termed dynamic instability, is based on GTP hydrolysis of the β-tubulin subunit which poises filaments for disassembly. It also allows microtubules to efficiently explore the intracellular space and generate forces on cellular structures such as kinetochores and the cell cortex (Brouhard and Rice, 2018). The cellular functions of microtubules are controlled by microtubule-associated proteins (MAPs) which serve to regulate microtubule dynamics and organize microtubules into supramolecular structures such as the mitotic spindle. MAPs appear to have co-evolved with tubulin and are already present in the last eukaryotic common eukaryotic ancestor (LECA) to regulate microtubule-dependent functions in the cell (Wickstead et al., 2010). How tubulin polymers mediate DNA segregation and cell division has mostly been studied in eukaryotes but homologous proteins are also present in bacterial phages and archaea (Fink and Lowe, 2015; Yutin and Koonin, 2012).

EB proteins are unique and particularly important plus end-interacting proteins (+TIPs) in eukaryotic cells (Akhmanova and Steinmetz, 2008; Schwartz et al., 1997; Tirnauer et al., 2002). Pioneering *in vitro* reconstitution experiments have revealed that EBs can localize to polymerizing plus-ends autonomously, i.e. in absence of proteins other than tubulin (Bieling et al., 2007). They accomplish this by recognizing structural features unique to the GTP hydrolysis state of the microtubule, resulting in an increased residence time of EB molecules at the plus-end versus the microtubule lattice (Maurer et al., 2012). Taking advantage of this property, EBs can recruit and enrich other proteins at plus-ends (Akhmanova and Steinmetz, 2008). Binding to EBs is established through protein-protein interactions between the EB-homology domain (EBH domain) and short linear interaction motifs (SLIMs) in the partner proteins (Honnappa et al., 2009; Kumar et al., 2017). Widespread EB binding motifs are SxIP, LxxPTPh, or combinations of these motifs (Buey et al., 2012; Jiang et al., 2012; Stangier et al., 2018). Sometimes these motifs generate a sufficiently high binding affinity towards EBs which can enrich cargo proteins at plus-ends. It remains difficult to predict which protein may be a physiologically relevant binding partner of EBs in cells, as potential EB interaction motifs are present in many different proteins, generate only modest affinity to EBs and need to compete with many other proteins for a limited number of EB molecules in the cell. Moreover, extensive synthetic genetic interaction studies between deletion mutants have implicated Bim1, the only yeast EB protein, in a broad range of cellular processes (Costanzo et al., 2016; Tong et al., 2001). The phenotypes of the respective double mutants, however, have typically not been characterized in detail, making it difficult to assess the precise contribution of Bim1 to these processes.

Among the EB-interacting proteins, the CLIP-170 family of proteins (Bik1 in budding yeast), plays a distinct role. It uses a different binding mode to interact with EBs, via an N-terminal CAP-Gly domain which interacts with the carboxy-terminal aromatic residues of EBs (Stangier et al., 2018). This interaction is sufficient to recruit CLIP-170 proteins to growing microtubule plus-ends (Bieling et al., 2008). Because of this binding mode, Bik1 can participate in Bim1 complexes with other proteins and potentially modulate their functions (Moore et al., 2006). It is, however, unclear, which functions are executed by Bim1-Bik1 complexes in the cell, compared to functions that can be fulfilled by either partner independently of each other.

Here we take advantage of the reduced complexity of the microtubule network in *S. cerevisiae* and the fact that this organism expresses only a single EB protein. As *BIM1* is a non-essential gene, the analysis of a microtubule cytoskeleton in the complete absence of an EB protein is possible. In addition, Bim1 mutants, specifically defective in different types of interactions with cargoes and Bik1, can be generated and analyzed. In this study, we set out to answer the following questions: Which microtubule-based processes are primarily affected by Bim1? What are the key Bim1-interaction partners in these processes and which strategies do cells employ to organize the microtubule cytoskeleton in the absence of an EB protein? Our results indicate that artificial plus-end targeting of a single cargo Kinesin-14 Cik1-Kar3 is sufficient to rescue most spindle-related *bim1* deletion defects in yeast cells. Upregulation of the Dynein-Dynactin nuclear migration pathway by increased accumulation of Bik1-Kip2 on the lattice of long astral microtubules compensates for the absence of the Bim1-Kar9-Myo2 complex in spindle orientation. Our experiments define a minimal set of Bim1 cargoes required to build and position a spindle for successful chromosome segregation.

## Results

### A microscopy-based screen for Bim1-dependent microtubule localization in yeast cells

To fully define the contribution of Bim1 to cellular functions in yeast, we aimed to systematically study the consequences of a *bim1* deletion for the localization of a wide variety of proteins implicated in microtubule-related processes. To this end we assembled a collection of proteins based on the following criteria: 1) reported involvement in the yeast spindle orientation pathways or nuclear spindle functions, 2) reported association with Bim1-Flag in IP-MS (van der Vaart et al., 2017), and 3) predicted SxIP motifs, or other putative Bim1 binding domains in unstructured regions, In addition, we confirmed that C-terminal tagging did not compromise the function of the tested proteins (**Supplementary Figure 1, Supplementary Table 1**). In total we tested 21 GFP or 3xGFP fusion proteins. To avoid cellular adaptation and accumulation of suppressor mutations, we constructed the GFP fusions in a hemizygous *bim1Δ* strain and imaged pairs of wild-type and *bim1Δ* mutants shortly after dissecting the diploids (**Figure 1A**). All GFP-fusions were expressed to similar levels in wild-type and *bim1* deletion strains (**Supplementary Figure 1B**). Live cell imaging was performed under standardized conditions for all strains following α-factor release from a G1-like state. The fluorescence intensity of the respective GFP-fusion was quantified along the spindle, or along a bud-directed cytoplasmic microtubule, respectively. The spindle pole body marker Spc42-mCherry was used to follow spindle morphogenesis (**Figure 1A**). Out of 21 GFP-fusions only two proteins fully depended on Bim1 for association with microtubules under these conditions (**Figure 1B**). The protein Kar9 in the cytoplasm, and the Kar3 (Kinesin-14) partner protein Cik1 in the nucleus. Cik1 association with the spindle was nearly abolished in a *bim1Δ* mutant (**Figure 1C**), consistent with the finding that binding motifs in the N-terminus of Cik1 and Kar3 are necessary for the formation of a ternary Bim1-Cik1-Kar3 complex (Kornakov et al., 2020). Spindle association of Kar3 itself was reduced, but not fully eliminated in the *bim1Δ* mutant, consistent with the formation of an alternative Kar3 motor complex with the Cik1 paralog Vik1, which localizes in a Bim1-independent manner to the spindle pole bodies (Manning et al., 1999; Mieck et al., 2015).

**Figure 1.**
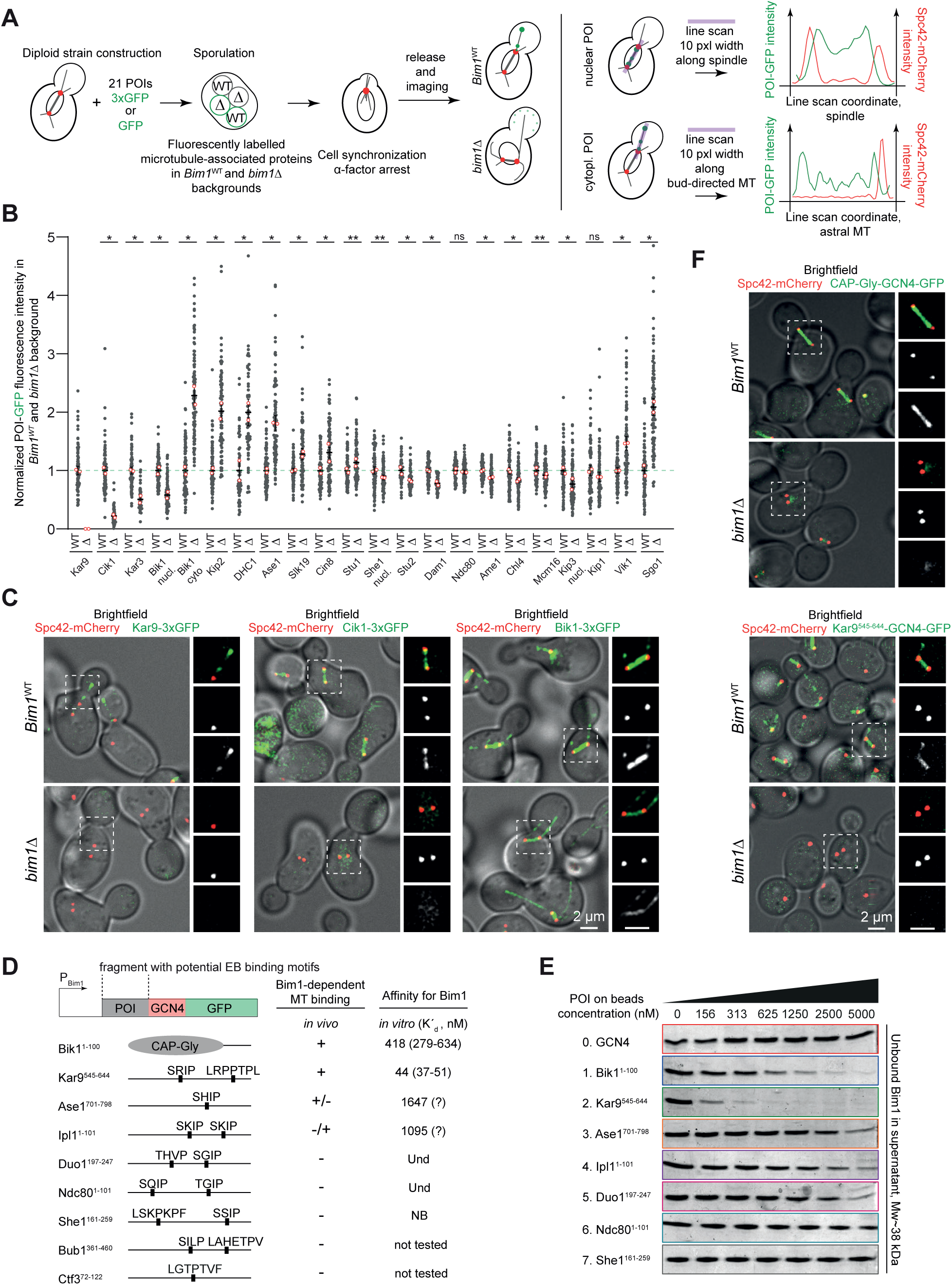
A microscopy-based screen reveals the effects of a *bim1* deletion on a collection of microtubule-associated proteins. **(A)** Schematic overview of strain construction and quantitative imaging of selected proteins in *bim1* deletion cells (see details in Methods chapter). **(B)** Overview of changes in microtubule localization to spindle or cytoplasmic microtubules of selected MAPs, fluorescence intensity of the respective MAP was normalized to one in the Bim1 wild-type strain. Small dots correspond to individual cells, open red circles represent mean values from two repeats. Error bars are mean values with 95% confidence intervals (CIs). More than 50 cells from two repeats were quantified for each condition. Asterisks denote: **, P≤0.0001; *,P≤0.01; ns, non-significant; unpaired two-tailed Student’s *t* test was used. **(C)** Selected examples for effects of *bim1*Δ on microtubule-associated proteins. Left side: Overlay of fluorescence microscopy with bright field image. On the right side of every image magnification of the boxed area is shown. From top to bottom: Magnification of boxed area, merged image, red channel, green channel. Scale bar, 2 μm. **(D)** Test for sufficiency of proposed Bim1 binding regions on the localization to microtubules *in vivo* and Bim1 binding *in vitro*. Selected fragments were cloned upstream of a GCN4 leucine zipper fused to GFP. For *in vivo* binding: +, localization to the spindle in all cells; +/-, most cells have signal above background; -/+, a few cells have detectable signal; -, no localization observed. More than 100 cells from two repeats were examined. For *in vitro* binding a dissociation constant with 95% CI is shown calculated from three repeats. Labels: Und, undefined, for constructs whose dissociation constant is outside of the tested experimental range; NB, no binding. **(E)** Pull-down assay with immobilized GCN4 fusion constructs. Coomassie-stained samples showing Bim1 in the supernatant fractions of the pull-down. **(F)** Examples of GCN4-GFP fusion proteins to the Bik1-, or Kar9-Bim1 interaction domains expressed from a *BIM1* promoter in wild-type or *bim1Δ* cells. Organization of the microscopy images as in C. Scale bar, 2 μm.

The deletion of Bim1 had differential effects on other spindle proteins: While the overall level of Bik1/CLIP-170 on the metaphase spindle was reduced by half, the association of other proteins, in particular the conserved crosslinking protein Ase1/PRC1, displayed a two-fold increase (**Figure 1B, Supplementary Figure 1C**). The same trend was observed for other MT crosslinkers such as Slk19, the Kinesin-5 family member Cin8 and the budding yeast CLASP homolog Stu1, which all showed a small but significant increase. For the kinetochore proteins tested in our collection, a slight reduction in their levels were observed for all subcomplexes, besides the major microtubule binding component of the kinetochore Ndc80 (**Figure 1B, Supplementary Figure 1D**). Robustness of Ndc80 loading might be achieved by the coexistence of multiple kinetochore assembly pathways or alternatively determined by intrinsic Ndc80 properties (Schleiffer et al., 2012). Reduction of Dam1c at the kinetochore upon Bim1 deletion was observed recently and has been attributed to the direct interaction between Duo1 and Bim1 (Dudziak et al., 2021).

Upon analyzing the associations in the cytoplasm, we found that Kar9-3xGFP foci on bud-directed cytoplasmic microtubules were abolished in the *bim1Δ* strain, consistent with earlier reports (Miller et al., 2000). Interestingly, the *bim1Δ* mutant had opposite effects on the microtubule-association of Bik1-3xGFP in the cytoplasm versus in the nucleus: While the spindle association of Bik1 was reduced, the *bim1Δ* strain displayed a two-fold increased level of Bik1 decorating long cytoplasmic microtubules **(Figure 1B, C)**. Bik1 appeared to be less confined to plus-ends, but instead could be detected in foci along the lattice, in line with previous observations (Stangier et al., 2018). Along with Bik1, the plus-end directed Kinesin Kip2 (Carvalho et al., 2004) was also strongly enriched on cytoplasmic microtubules in the *bim1Δ* mutant. We conclude that only Kar9 and Cik1 strictly rely on Bim1 for microtubule association in cells and that a major effect of the *bim1Δ* mutant is an apparent redistribution of the CLIP-170 homolog Bik1 from the nucleus into the cytoplasm.

### Fluorescent reporter constructs recapitulate Bim1-dependent binding in cells

The strict dependence of Kar9 and Cik1-Kar3 on the presence of Bim1, as well as the different effects of *bim1Δ* on nuclear and cytoplasmic Bik1, may reflect the formation of stable complexes between Bim1 and these binding partners in cells. To study whether Bim1 interaction motifs on selected candidate proteins are sufficient to recapitulate the localization of Bim1 cargoes in cells, we prepared constructs containing the predicted Bim1-binding regions dimerized by the Leucine-Zipper domain of GCN4 and fused to GFP (**Figure 1D**). We then asked if these constructs could directly interact with recombinant Bim1 *in vitro*. In parallel we analyzed their localization in cells. Using quantitative pull-down assays, we found that the different constructs interacted with Bim1 with binding affinities decreasing in the order Kar9>Bik1>Ipl1≃Ase1>>others (**Figure 1D, E**, **Supplementary Figure 1E**). The CAP-Gly domain construct, expressed from a *BIM1* promoter, almost exclusively localized to the spindle of yeast cells. This localization was dependent on Bim1 (**Figure 1F**). This implies that other binding partners of the CAP-Gly domain, such as the carboxy-termini of α-Tubulin (Tub1 and Tub3), are insufficient to localize this construct to the spindle or to cytoplasmic microtubules. We speculate that the CAP-Gly domain of yeast Bik1 has a much higher affinity towards Bim1 (-ETF*) than to tubulin tails (-EEF*) within microtubules or, perhaps less likely, that tubulin tails are pre-blocked by other proteins in the cell. The Kar9 construct, on the other hand, localized to both cytoplasmic and spindle microtubules, and no microtubule association could be observed in a *bim1* deletion mutant (**Figure 1F**). Ase1 and Ipl1 constructs, which bind Bim1 with low micromolar affinity *in vitro* were poorly recruited to spindles *in vivo*. Thus, strict dependence on Bim1 in cells correlates with a high affinity for Bim1-binding motifs *in vitro*.

### A *bim1* deletion alters the spindle association dynamics of several microtubule-associated proteins

We next characterized the intranuclear phenotypes of *bim1Δ* cells by imaging spindle assembly in synchronized cells with 2 minutes time resolution. To visualize Bim1 itself, we constructed a new Bim1-GFP fusion (termed Bim1^WT^-GFP) by inserting GFP with short linker sequences into the unstructured region between the EBH domain and the extreme C-terminus (**Figure 2A**). This construct avoids blocking the N-or C-terminus and in particular allows CAP-Gly domain-mediated binding of Bik1 to the C-terminal - ETF* motif of Bim1 (Stangier et al., 2018). The construct fully rescued the temperature-, hydroxyurea- and benomyl-hypersensitivities of a *bim1Δ* mutant. In addition, Bim1^WT^-GFP was able to support cell viability in the *mad1Δ* background upon deletion of the spindle assembly checkpoint (SAC) indicating error-free chromosome segregation (**Supplementary Figure 2A**). Bim1^WT^-GFP displayed a prominent spindle localization and a weak signal at the ends of astral microtubules. At both locations Bim1 co-localized with Bik1 and on the spindle it co-localized with Stu2 (**Supplementary Figure 2B**). Using high time resolution microscopy we examined Bim1 behavior on dynamic microtubules in α-factor arrested cells. Bim1 tracked polymerizing and depolymerizing ends of cytoplasmic microtubule bundles as well as individual microtubules aligned along the bundles. Depolymerizing ends of individual microtubules were weakly stained with Bim1 (**Supplementary Figure 2C**). We then compared the association dynamics of Bim1 itself with that of other spindle MAPs: Bik1-3xGFP and Bim1^WT^-GFP spindle association dynamics closely mirrored each other. Both proteins accumulated early on metaphase spindles, reached peak levels in the middle of metaphase and declined upon anaphase onset (**Figure 2B, D**, **Supplementary Video 1**). The remaining weak Bim1 and Bik1 signals in anaphase were localized at the spindle midzone, at kinetochore clusters and at individual dots along the spindle (**Supplementary Figure 2B**). Consistent with the relatively low levels of Bim1 on anaphase spindles, parameters such as maximal anaphase spindle length, as well as anaphase spindle elongation kinetics were largely unaffected in *bim1* deletion cells under our imaging conditions (**Figure 2C, E**). In contrast to Bim1-Bik1, the level of the CLASP protein Stu1-GFP was relatively constant over time. The protein was present both on metaphase and anaphase spindles until spindle disassembly (**Figure 2B, D**). Ase1-GFP was initially absent on the nascent metaphase spindle, its level then gradually increased until a maximum was reached shortly before anaphase spindle disassembly. Wild-type cells had 0.92±0.03 μm initial metaphase spindle length which gradually increased to 1.98±0.04 μm before anaphase onset (mean ± SEM). Cells lacking *BIM1* were characterized by a shorter initial and maximal metaphase spindle length, 0.67±0.01 μm and 1.48±0.05 μm, respectively (**Figure 2C, E**). Moreover, we found that metaphase spindle length typically increased monotonously in wild-type cells, whereas *bim1Δ* cells displayed more frequent fluctuations of metaphase spindle length over time (**Supplementary Figure 2D**). Abrupt changes in metaphase spindle length of *bim1Δ* cells were often associated with transient pulling and pushing events, when the spindle was tumbling between mother cell and bud (**Supplementary Video 1 and 2**). A similar behavior has previously been reported for *ase1Δ* and could be explained by dynein-dependent forces applied on astral microtubules connected to a spindle that is compromised in microtubule crosslinking (Estrem and Moore, 2019). Since we established that *bim1Δ* cells loaded much more Ase1 on their short spindles, we decided to investigate the timing of Ase1 recruitment. Surprisingly, we found that in *bim1Δ* cells Ase1 was already present at a detectable, albeit low, level before bi-polar spindle formation (**Figure 2C, Supplementary Video 1**). As mitosis progressed, the Ase1 signal continuously increased and was higher than in wild-type cells during both metaphase and anaphase (**Figure 2C, D**).

**Figure 2.**
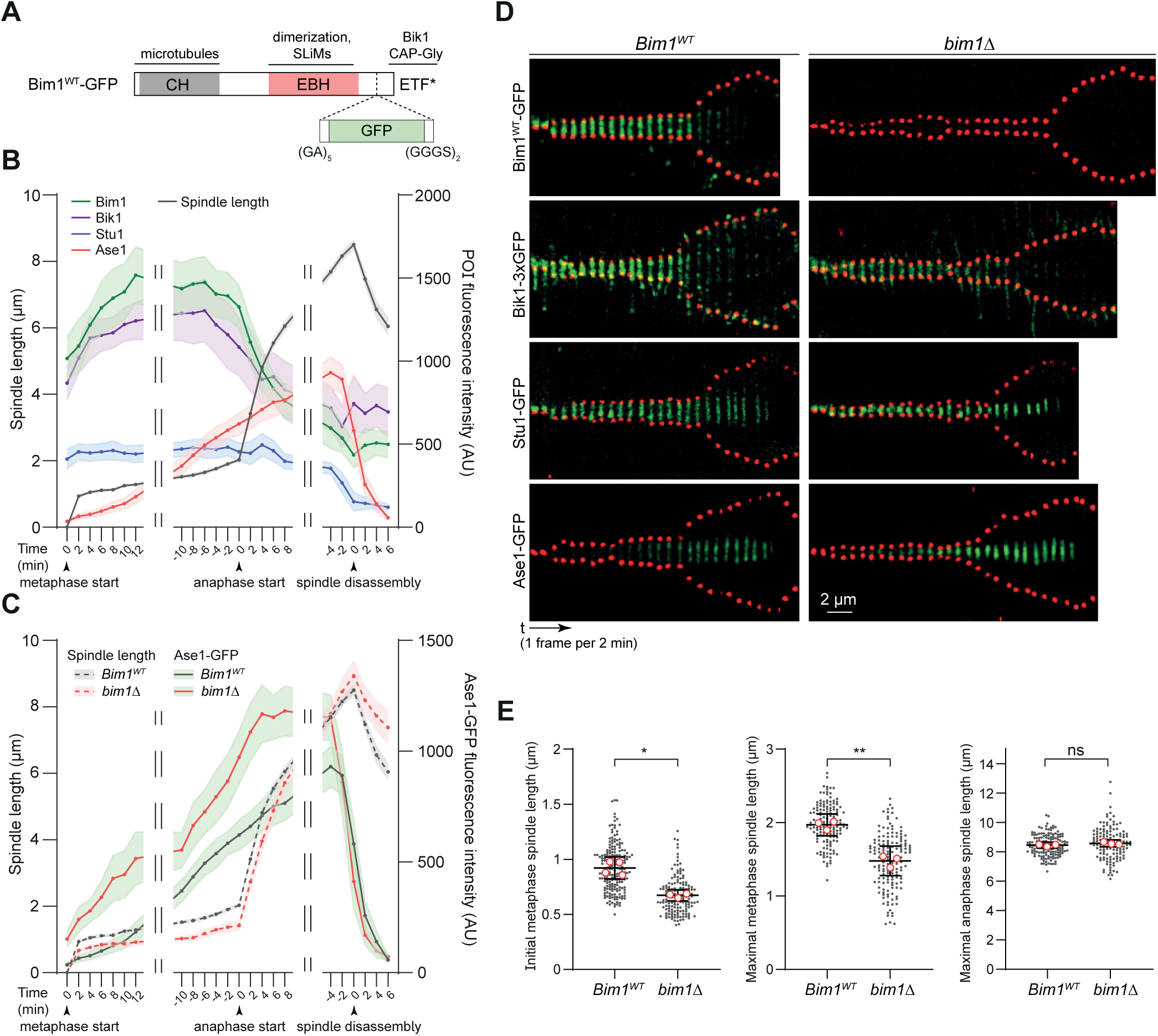
Spindle association dynamics of several MAPs is altered in *bim1* deletion cells. **(A)** Scheme for the construction of a functional Bim1-GFP fusion protein. GFP is inserted between the EBH domain and extreme C-terminus. **(B)** Quantification of fluorescence intensity and spindle length as a function of time in wild-type cells. Start of metaphase, start of anaphase and start of spindle disassembly is indicated in the graph, time (in minutes) is given relative to these reference points. Breaks in the graph are used to show correction for differences in metaphase and anaphase durations. Bim1, Bik1, Stu1 and Ase1 were quantified during spindle morphogenesis, beginning immediately after spindle pole body duplication until the start of disassembly of the anaphase spindle. For spindle size 75 cells were analyzed. For fluorescence intensity 20 Bim1-GFP, 15 Bik1-3xGFP, 15 Stu1-GFP, 25 Ase1-GFP cells were quantified. Curves show mean values with 95% CIs. (**C)** Quantification of fluorescence intensity of Ase1-GFP (solid lines) and spindle length (dotted lines) over time on wild-type or *bim1Δ* spindles. Data for wild-type background is the same as in **(B)**. In the *bim1Δ* background 25 cells were analyzed for spindle size and 15 cells for Ase1-GFP fluorescence intensity. **(D)** Representative reslices of selected GFP-fusions throughout metaphase and anaphase of mitotic spindles in wild-type or *bim1Δ* cells. Images were scaled to equal intensity between wild-type and *bim111*. Scale bar 2 μm, 1 frame/2 min. (**E)** Quantification of initial metaphase spindle length, maximal metaphase spindle length and maximal anaphase spindle length for Bim1 wild-type and *bim1Δ* cells (N cells were analyzed in every experiment: 200 and 150; 150 and 150; 150 and 135). Small dots correspond to individual cells, open red circles represent mean values of biological repeats. Error bars are mean values with 95% CIs. Asterisks denote *, P=0.0014; *,P=0.0013; ns, non-significant (p=0.1551); unpaired two-tailed Student’s *t* test was used.

In addition to Ase1, we followed the kinetochore proteins Ndc80-GFP and Sgo1-GFP which specifically marks kinetochores that lack tension (Indjeian et al., 2005). We observed that *bim1Δ* cells had mispositioned kinetochores with a bright Sgo1-GFP signal that was much stronger than in wild-type cells (**Supplementary Figure 2E**, **Figure 1B**). Lastly, we examined the Kinesin-5 Cin8-3xGFP as the earliest microtubule crosslinker driving SPB separation (Leary et al., 2019). Localization of Cin8-3xGFP was overall similar between wild-type and *bim1Δ* cells, however in the latter it was more confined at kinetochore clusters (**Supplementary Figure 2E**). Our observations are consistent with the idea that ordered recruitment of microtubule crosslinkers to the spindle reflects distinct functional contributions (**Supplementary Figure 2F**). We speculate that Bim1-Bik1 in a complex with its cargo Cik1-Kar3 is active after bi-polar spindle formation but before late metaphase and Ase1 can partially substitute for nuclear Bim1 functions.

### Bim1 confers its major function for the metaphase spindle via the kinesin-14 motor Cik1-Kar3

Further we compared the *bim1Δ* phenotype to deletions of its main cargoes Kar3 and Kar9. Analyzing spindle elongation kinetics in the respective deletion strains showed that a *kar3* deletion indeed closely reflects the *bim1* deletion. *kar3Δ* spindles were initially shorter, had a decreased maximum metaphase spindle length while metaphase duration increased to 47±2 min (48±2 min for *bim1Δ* and 31±1 min in wild-type cells) (**Figure 3A, B, C**). Unexpectedly, the *kar9* deletion mutant displayed a slightly accelerated metaphase progression relative to wild-type cells (26±1 min) (**Figure 3C**). This could be attributed to an increased level of Bim1 on the metaphase spindle of *kar9Δ* (or *Kar9-AID*) cells (**Figure 3D, E**). In this context we noticed that *kar9* deleted cells, as well as cells acutely depleted of Kar9 (*Kar9-AID*) almost completely lack cytoplasmic Bim1, while the overall level of Bim1 in western blots was unchanged (**Figure 3D, F, G**). Thus, Kar9 and Bim1 are mutually dependent for their cytoplasmic localization and lack of Kar9 impacts the distribution of Bim1 between the cytoplasm and the nucleus. This finding supports a recently published model which states that nuclear-cytoplasmic shuttling of Kar9 controls the cytoplasmic localization of Bim1 (Schweiggert et al., 2016). Cik1 depletion had no detectable effect on the amount of Bim1 on the spindle, but instead caused an increased recruitment of Bim1 on astral microtubules (**Figure 3D, E, F**). This might serve as an indication that Kinesin-14 contributes to the nuclear import of Bim1 during mitosis. Loading of Bim1 on the spindle was partially dependent on Bik1. Bik1 depletion led to a coordinated decrease in nuclear and increase in cytoplasmic Bim1 and cytoplasmic microtubules were very short in the absence of Bik1 (**Figure 3D, E, F**). Overall, our cell biology data suggested that major nuclear Bim1 functions are conducted in a complex with Cik1-Kar3, while Bik1 and Kar9 have a smaller impact, probably affecting the nuclear-cytoplasmic distribution of Bim1.

**Figure 3.**
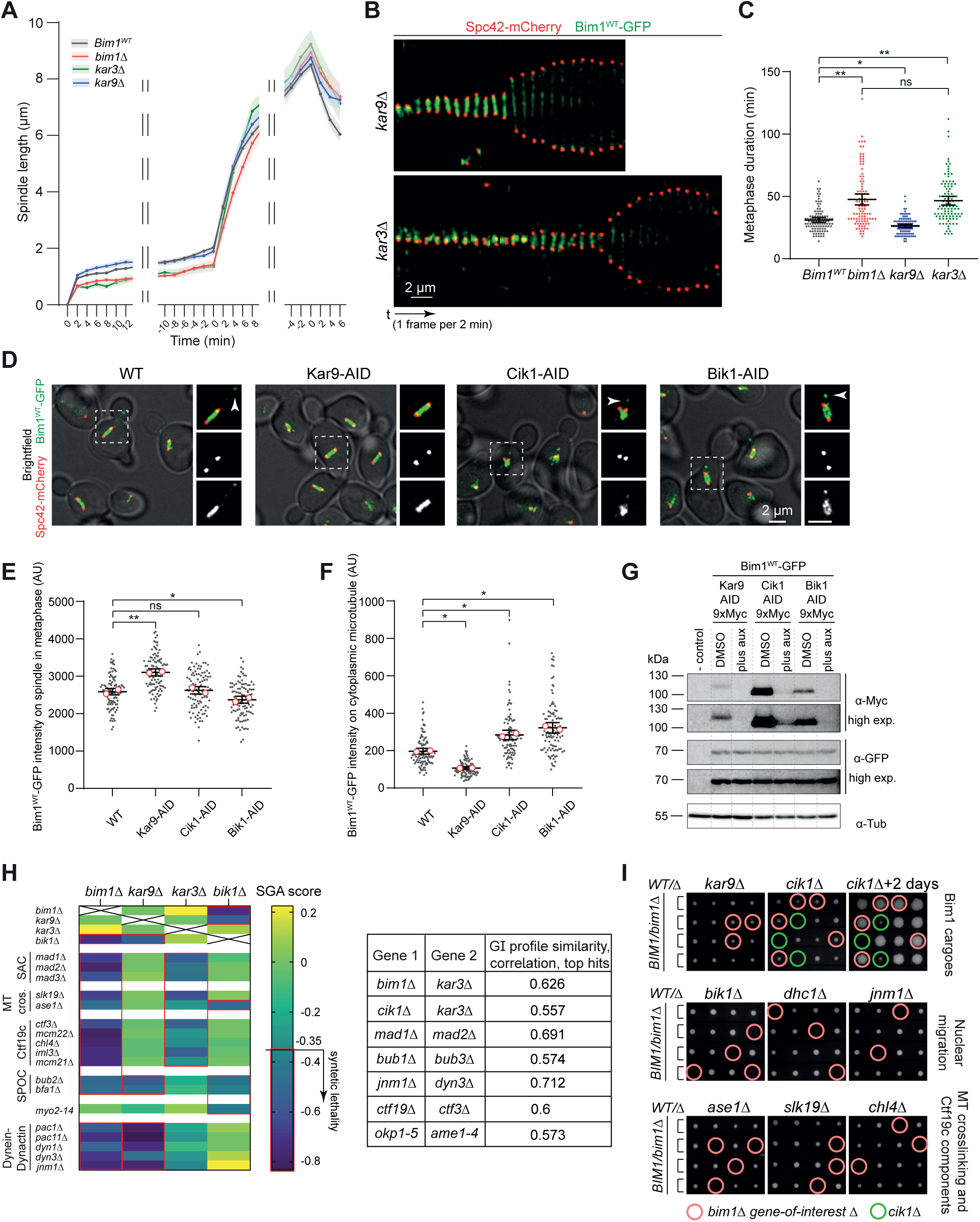
Bim1 executes its main nuclear function via the Kinesin-14 Kar3. **(A)** Quantification of metaphase and anaphase spindle elongation kinetics in wild-type, *bim1Δ, kar9Δ or kar3Δ cells*. 25 spindles were analyzed for *kar9Δ or kar3Δ cells* (data for wild-type and *bim1Δ* cells is the same as in Figure 2C). Curves show mean values with 95% CIs. Note that metaphase spindle elongation profiles for *bim1Δ* and *kar3Δ* mutants closely resemble each other. **(B)** Kymographs of Bim1 association during metaphase and anaphase on spindles in a *kar9* and a *kar3* deletion. Note reduced metaphase spindle size in the *kar3Δ* strain. Scale bar 2 μm, 1 frame per 2 min. **(C)** Quantification of metaphase duration in the indicated strains. For each condition 100 cells were quantified. Error bars are mean values with 95% CIs. Asterisks denote **, P<0.0001; *,P=0.0005; ns, non-significant (p=0.9841); Welch and Brown-Forsythe ANOVA with Games and Howell post-test was used. **(D)** Effects of acute depletion of Kar9, Cik1 or Bik1 on the localization and distribution of Bim1. Fluorescence micrographs showing Bim1^WT^-GFP and Spc42-mCherry in wild-type cells or upon acute depletions of Kar9, Cik1 or Bik1. Left side: Overlay of fluorescence microscopy with bright field image. On the right side of every image magnification of the boxed area is shown. From top to bottom: Magnification of boxed area, merged image, red channel, green channel. Arrowheads point to Bim1^WT^-GFP on cytoplasmic microtubules. Scale bar, 2 μm. **(E)** Quantification of Bim1^WT^-GFP intensity on the metaphase spindle in wild-type or depletion strains. Note increased level of Bim1 on metaphase spindle upon depletion of Kar9. 100 of cells were analyzed for each condition. Error bars are mean values with 95% CIs. Asterisks denote: **, P<0.0001; *,P=0.0019; ns, non-significant (p=0.9517); Welch and Brown-Forsythe ANOVA with Games and Howell post-test was used. **(F)** Quantification of Bim1^WT^-GFP intensity on the cytoplasmic microtubules in wild-type or depletion strains. 100 of cells were analyzed for each condition. Error bars are mean values with 95% CIs. Asterisks denote *, P<0.0001; Welch and Brown-Forsythe ANOVA with Games and Howell post-test was used. **(G)** Western blot showing effectiveness of depletion of Kar9, Cik1 and Bik1. Whole cell extracts were separated by SDS-PAGE, blotted and probed with anti-myc (AID-fusion) or anti-GFP antibody (Bim1). To enhance the Cik1 signal, α-factor arrested cells were used which express elevated levels of Cik1. **(H)** Left side: Synthetic genetic analysis assigning SGA scores to combinations of deletion mutants. SAC, spindle assembly checkpoint; SPOC, spindle position checkpoint. Synthetically lethal interactions are concluded in the red box. On the right side Pearson correlation is shown for SGI profiles of selected genes. **(I)** Heterozygous diploid dissection of different gene deletions in a *bim1Δ* background. Four spores of a tetrad are displayed horizontally; the indicated genotypes are marked by circles. Note that *bim1Δ kar9Δ* and *bim1Δ cik1Δ* double mutants are viable.

Next, we compared the comparing genetic interaction profile of a *bim1* deletion to that of various other factors by re-analyzing the synthetic genetic interaction data from (Costanzo et al., 2016) and TheCellMap.org (Usaj et al., 2017). A comparison of the genetic interaction profiles shows that a *kar3* deletion was most similar to a *bim1* deletion, while a *kar9* deletion resembled the *bim1* deletion profile only with regards to mutants in the Dynein-Dynactin complex (**Figure 3H, Supplementary Table 2**). Quantitative correlation between genetic interaction profiles of *bim1Δ* and *kar3Δ* was in the same range as for members of established multi-protein complexes, such as Bub1-Bub3, Mad1-Mad2 or the centromere-associated proteins Ctf19 and Ctf3 (**Figure 3H**). If Bim1 confers its functions indeed in the form of Bim1-Cik1-Kar3 complexes in the nucleus and Bim1-Kar9 complexes in the cytoplasm, then combinations of the respective deletion mutants should not show synthetic defects, but might display epistasis (van Leeuwen et al., 2016). Indeed, *bim1Δ kar9Δ* and *bim1Δ cik1Δ* double mutants were viable and did not show aggravated phenotypes relative to the individual mutants (**Figure 3I**). The *bim1* deletion even improved the growth of *cik1Δ* spores. In contrast, *bim1Δ* was synthetic lethal with mutants involved in nuclear migration, microtubule crosslinking, tension sensing and centromeric cohesion (**Figure 3H, I**). Genetic analysis suggested a primary role of Bik1 in the Dynein-Dynactin nuclear migration pathway that would work in parallel to Bim1-Kar9-Myo2. Bim1-Cik1-Kar3 in turn contributes to the assembly of a proper spindle structure and chromosome bi-orientation in parallel to Ase1-Slk19. Since we have found that *bim1Δ* leads to accumulation of Bik1 on cytoplasmic microtubules, we decided to analyze the cytoplasmic events of spindle positioning and nuclear migration in more detail.

### Anaphase spindle elongation occurs in the absence of Bim1 and Bik1, but the nucleus fails to migrate into the bud neck

We sought to define the terminal phenotypes of mutants simultaneously lacking Bim1 and either the microtubule crosslinker Ase1 or the CAP-Gly protein Bik1, as deletions of both proteins were synthetic lethal with *bim1Δ*. To this end we combined the *bim1* deletion mutant with auxin-inducible degron alleles of either Bik1 or Ase1. The cells expressed Sgo1-GFP as a marker for tension establishment and Spc42-mCherry to monitor spindle size and position. Cells were imaged after release from an α-factor arrest into auxin-containing medium. (**Figure 4A**). Wild-type cells quickly proceeded through metaphase, reached a 1.61±0.02 μm average spindle length, cleared off Sgo1-GFP as cells achieved bi-orientation, positioned the spindle in the bud neck and elongated it between mother and bud in anaphase (**Figure 4A, B, C, D**). Depletion of Bik1 resulted in a mild decrease of metaphase spindle size (1.39±0.02 μm) but neither a delayed metaphase progression nor an activated tension checkpoint. Cells positioned the spindle properly but underwent spindle elongation almost exclusively within the mother cell. They only managed to conclude nuclear migration during a delay in anaphase (**Figure 4A, D**). Ase1 depletion led to a strong decrease in metaphase spindle size similar to *bim1Δ* (1.11±0.02 μm and 1.05±0.02 μm, respectively), but in contrast to *bim1Δ* did not cause accumulation of Sgo1-GFP and had no metaphase delay. Anaphase spindle elongation was severely compromised in Ase1 depleted cells and about ∼40% of cells initially performed this in the mother cell, only later pulling the spindle through the bud neck. The combinations of the *bim1Δ* allele with the two different degrons showed distinct phenotypes: Many *Bik1-AID bim1Δ* cells managed to clear Sgo1-GFP from the spindle and initiated anaphase chromosome segregation. Spindle elongation in these cells, however, occurred nearly exclusively in the mother cell (**Figure 4A, D**). The spindles of *Ase1-AID bim1Δ* cells, on the other hand, remained very short (0.83±0.02 μm) and were prominently decorated with Sgo1-GFP; these cells did not initiate anaphase during the course of the experiment (**Figure 4A, B**). Thus, both double mutants eventually fail to distribute chromosomes between mother and daughter cells, but do so with distinctly different terminal phenotypes. Overall, our live cell analysis is in a good agreement with the genetic data and indicates the existence of two parallel and partially redundant pathways for spindle assembly and chromosome bi-orientation (Bim1-Cik1-Kar3 and Ase1-Slk19) and for spindle positioning and nuclear migration (Bim1-Kar9-Myo2 and Bik1-Kip2-Dynein-Dynactin). Since we previously found that *bim1Δ* leads to accumulation of Bik1 on cytoplasmic microtubules, we decided to analyze the cytoplasmic events of spindle positioning and nuclear migration in more detail.

**Figure 4.**
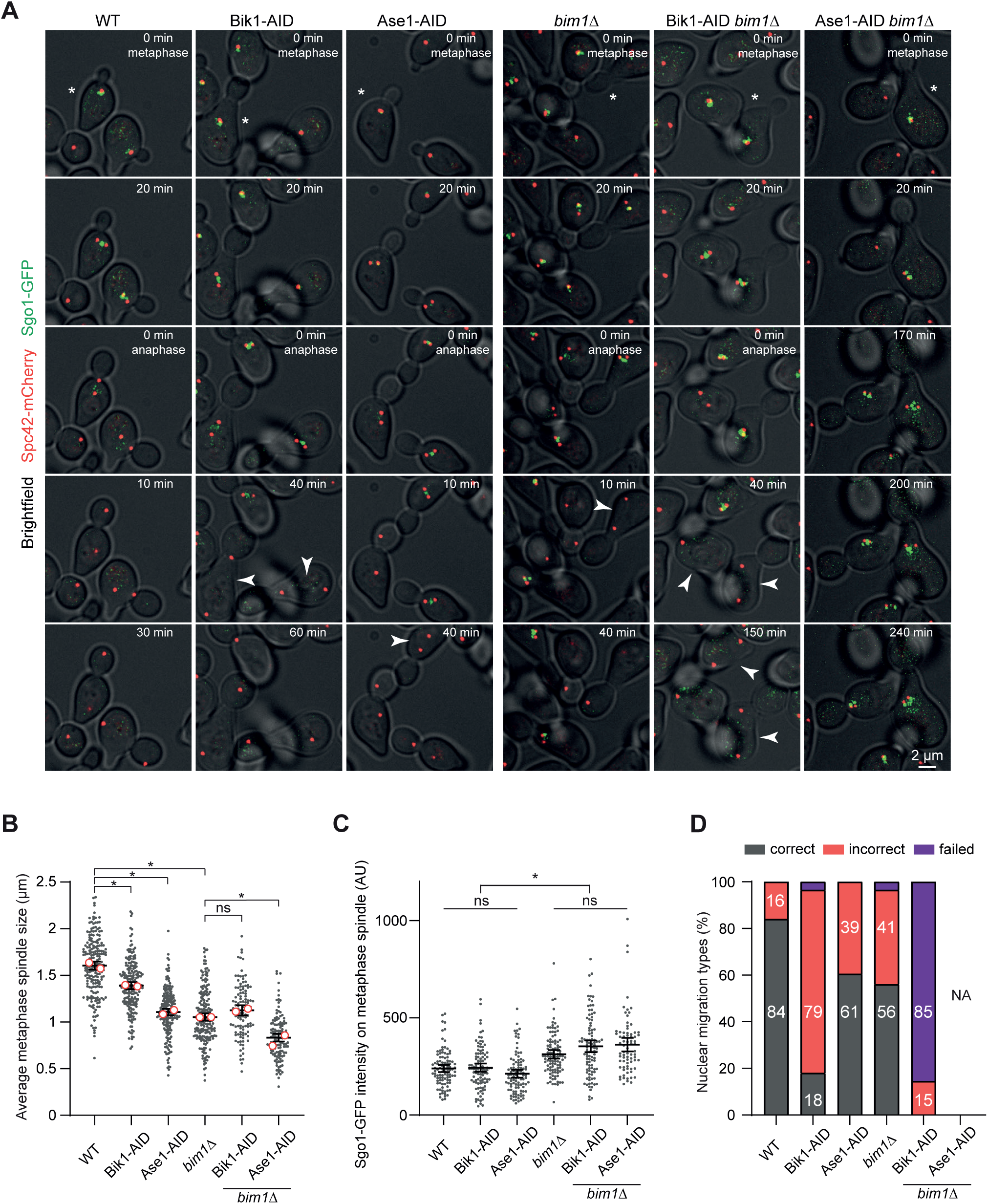
Distinct terminal phenotypes of *bim1Δ* cells lacking Ase1 or Bik1. **(A)** Live cell microscopy of cells expressing Sgo1-GFP and Spc42-mCherry in the indicated strain backgrounds. Cells were released from α-factor arrest into auxin-containing medium and imaged throughout the cell cycle (time lapse 10 min). Minutes indicate time into metaphase (after initial separation of SPBs) and time after anaphase onset (continued separation of spindle pole bodies) for the cell labelled with the star in the first frame. Arrowheads point at cells elongating anaphase spindles entirely in the mother cell. Scale bar 2 μm. **(B)** Quantification of average metaphase spindle size (20 min before anaphase) of the indicated strains. Number of cells analyzed for every condition: 200 wild-type, 200 *Bik1-AID*, 200 *Ase1-AID*, 200 *bim1Δ,* 110 *Bik1-AID bim1Δ*, 127 *Ase1-AID bim1Δ.* Small dots correspond to individual cells, open red circles represent mean values from two repeats. Error bars are mean values with 95% CIs. Asterisks denote *, P<0.0001; ns, non-significant (p=0.2534); Welch and Brown-Forsythe ANOVA with Games and Howell post-test was used. Note that *bim1Δ Ase1-AID* cells have shortened metaphase spindles relative to *bim1Δ* or *Ase1-AID* alone. **(C)** Quantification of Sgo1-GFP intensity on metaphase spindles (within 10 min after SPB separation) in the indicated strains. 100 cells were analyzed for each condition, with exception of 76 *Ase1-AID bim1Δ* cells. Small dots correspond to individual cells. Error bars are mean values with 95% CIs. Asterisks denote *, P<0.0001; ns, non-significant; Welch and Brown-Forsythe ANOVA with Games and Howell post-test was used. **(D)** Quantification of nuclear migration phenotypes. “incorrect” indicates initial anaphase spindle elongation in the mother cell, which is later corrected, “failed” indicates anaphase spindles that stay in the mother cell. NA: not applicable, since the phenotype is scored relative to anaphase onset and *Ase1-AID bim1Δ* cells do not progress past metaphase. Two repeats of 100 cells each were quantified, average values are shown.

### Spindle orientation and nuclear migration in the absence of Bim1 relies on an upregulation of the Dynein-Dynactin pathway via Bik1-Kip2

To characterize the cytoplasmic events that occur in the absence of Bim1, we measured the parameters of spindle orientation and nuclear migration in wild-type, *bim1!1* cells and in the deletions of its two main cargoes Kar9 and Kar3. To quantify the phenotypes we measured the distance between the spindle pole body and the bud-neck axis and evaluated nuclear migration across the bud neck two frames (4 min) after anaphase onset (**Figure 5A**). We found that *bim1Δ* and *kar9Δ* cells display spindle positioning defects, characterized by an increased distance between SPB and bud neck on pre-anaphase spindles (1.11±0.1 μm and 2.27±0.12 μm, compared to 0.58±0.05 μm in wild-type cells) (**Figure 5B, Supplementary Figure 5A**). Despite Bim1 and Kar9 localization on astral microtubules being co-dependent, surprisingly we found that both spindle orientation and nuclear migration defects were stronger in *kar9Δ* than in *bim1Δ* (**Figure 5B, C**). To better understand how *bim1* deletion cells accomplish spindle orientation and nuclear migration we further investigated components of the Dynein-Dynactin pathway in more detail. We measured that *bim1Δ* cells had longer astral microtubules than wild-type cells (4.28±0.12 μm and 2.05±0.07 μm correspondingly) (**Figure 5D, F**). Next, we asked how the amount of Bik1 in the cytoplasm would affect microtubule length. Bik1 is known to be required for astral microtubule stability and Bik1 in complex with its binding partner, the plus-end directed kinesin Kip2 (Carvalho et al., 2004), has microtubule polymerase activity *in vitro* (Hibbel et al., 2015) and *in vivo* (Chen et al., 2019). We constructed GFP-tagged Bik1 variants that were targeted to the nucleus or the cytoplasm via the inclusion of a nuclear localization signal (NLS) or a nuclear export signal (NES), respectively (**Figure 5D**). In agreement with our prediction that Bik1 is required in the cytoplasm, *Bik1-NLS* aggravated the growth defect of *bim1* deletion mutants (**Supplementary Figure 5B**). The strain displayed slow growth at 30°C and was nearly inviable at 37°C. By contrast the combination of *Bik1-NES* and *bim1Δ* displayed growth defects similar to the *bim1Δ* strain. Thus, Bik1 is specifically required in the cytoplasm of *bim1Δ* cells, where it contributes to the stability of cytoplasmic microtubules and participates in the Bik1-Kip2-Dynein-Dynactin complex. Targeting of Bik1 to the cytoplasm, however, was insufficient to increase microtubule length in wild-type or in *bim1Δ* cells. This makes a simple relationship between Bik1 accumulation and microtubule length unlikely (**Figure 5D, F**). The overall increase of Bik1 on astral microtubules in *bim1Δ* cells originated from the decoration of the lattice of long microtubules, despite the intensity of Bik1 at the plus-ends and in vicinity of the SPBs being reduced (**Figure 5E**, **Supplementary Figure 5C**). The majority of Bik1 foci on long microtubules displayed plus-end directed motility, comparable to the speed of Kip2 (∼6-12 μm/min) (**Figure 5G**). These observations are consistent with the idea that Bik1 acts as a processivity factor for Kip2: If more Bik1 is present on the lattice, then more Kip2 molecules are able to reach plus-ends without detachment (**Figure 5H**). Indeed, in contrast to Bik1, Kip2 displayed a prominent enrichment at the plus-ends in *bim1Δ* cells (**Figure 5I, J, Supplementary Figure 5D**). The increased levels of Bik1, Kip2, as well as the Dynein-Dynactin complex subunits Dhc1 and Jnm1, indicated an upregulation of this pathway in the *bim1* deletion cells (**Figure 5J, Figure 1B**). The hyperactive Dynein-Dynactin pulls on astral microtubules and promotes spindle tumbling in *bim1Δ* cells. In the absence of Bim1-Kar9-Myo2 this might be beneficial, since it reduces the distance from the spindle to the bud neck and increases chances for proper nuclear migration upon anaphase onset.

**Figure 5.**
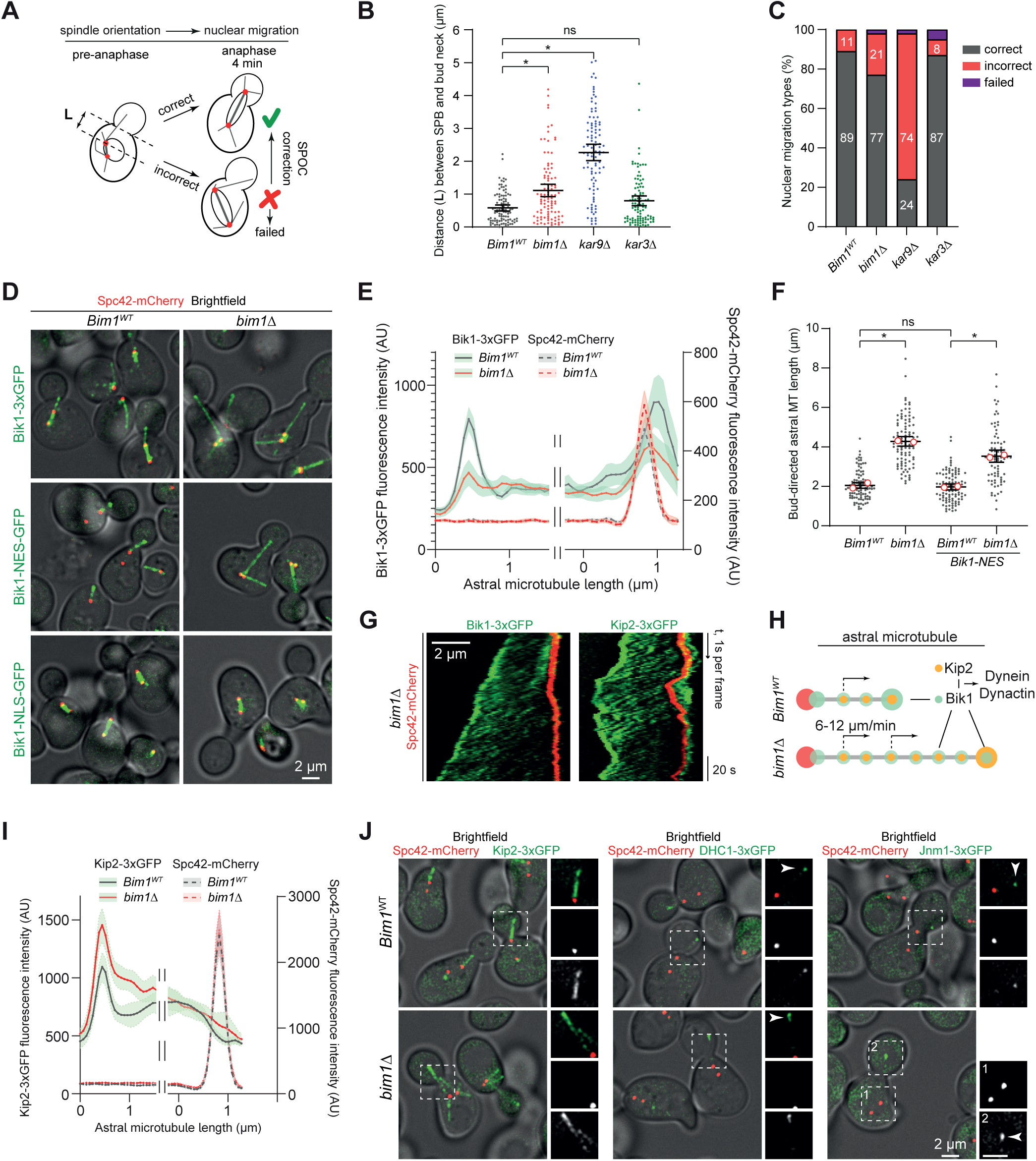
Nuclear migration in the absence of Bim1 requires cytoplasmic Bik1. **A)** Scheme for scoring spindle orientation and nuclear migration defects. **L** describes the distance between the bud-proximal spindle pole body and the bud-neck axis, nuclear migration was examined two frames (4 min) after anaphase spindle elongation. (**B)** Quantification of the distance between the bud-proximal spindle pole body and the bud-neck axis in different mutants. 100 cells were analyzed for each condition. Small dots correspond to individual cells. Error bars are mean values with 95% CIs. Asterisks denote *, P<0.0001; ns, non-significant (P=0.0866); Welch and Brown-Forsythe ANOVA with Games and Howell post-test was used. Note the difference between the severity of *bim1Δ* and *kar9Δ* phenotypes. **(C)** Nuclear migration categories of wild-type cells and different mutants. Two repeats of 100 cells each were quantified, average values are shown. Category “Failed” indicates that anaphase spindles remained completely in the mother cell. **(D)** Live cell microscopy of Bik1-3xGFP in wild-type form, or directed to Cytoplasm or Nucleus via inclusion of an NES or NLS, respectively. Imaging was performed in a wild-type or *bim1Δ* background. Spindle pole bodies are labelled with Spc42-mCherry. Scale bar, 2 μm. **(E)** Intensity profiles of Bik1-3xGFP along astral microtubules in wild-type or *bim1Δ* cells. Spc42-mCherry intensity profile is displayed for reference. Break in the graph corrects for different microtubule lengths. 30 wild-type and 25 *bim1Δ* cells were analyzed. Curves show mean values with 95% CIs. **(F)** Bud-directed astral microtubule length in wild-type and different mutants. 100 cells were analyzed for each condition and 80 *Bik1-NES bim1Δ* cells. Small dots correspond to individual cells, open red circles represent mean values from two repeats. Error bars are mean values with 95% CIs. Asterisks denote: *, P<0.0001; ns, non-significant (P=0.8923); Welch and Brown-Forsythe ANOVA with Games and Howell post-test was used. **(G)** Kymograph analysis of Bik1-3xGFP and Kip2-3xGFP on long cytoplasmic microtubules in *bim1Δ* strains. Scale bar, 2 μm; time-lapse 1s/frame. Note oblique lines indicating movement of Bik1 and Kip2. **(H)** Schematic representation of Kip2 and Bik1 on cytoplasmic microtubules. **(I)** Averaged intensity profiles of Kip2-3xGFP along astral microtubules in wild-type and *bim1Δ* cells. Spc42-Cherry profile is included for reference. 25 of wild-type and *bim1Δ* cells were analyzed. Curves show mean values with 95% CIs. Break in the graph corrects for different microtubule lengths. **(J)** Fluorescence micrographs of Dynein-Dynactin nuclear migration pathway members Kip2, Dynein heavy chain (DHC1) and Jnm1 in wild-type or *bim1Δ* cells. Overview merged with brightfield on the left side. Magnification of the boxed area, merge, red channel, green channel from top to bottom. Arrowheads point at individual Dhc1 or Jnm1 foci. Scale bar 2 μm.

### Spindle-related Bim1 functions can be provided efficiently by engineered plus-end targeting of Kinesin-14

Given the similarities between the *bim1* and the *kar3* deletion, we asked to what extent the nuclear phenotypes of the *bim1* deletion could be rescued by providing plus-end targeting of Kar3 in the absence of Bim1. To this end we fused the CH domain of Bim1 (residues 1-133) to full-length Cik1 and integrated this construct (expressed from a *CIK1* promoter) into *bim1* deletion cells. In essence, this operation generates a cell with only a single Bim1 cargo complex in the nucleus. We then characterized the phenotype of the derived stain. We found that in serial dilution assays the expression of *CH-Cik1* rescued the temperature-, hydroxyurea- and to a slightly lesser extent the benomyl-hypersensitivity of the *bim1* deletion (**Figure 6A**). Expression of *CH-Cik1* was sufficient to shorten the metaphase duration of *bim1Δ* cells back to wild-type timing (31±1 min, 45±1 min, 31±1 min for wild-type, *bim1Δ* and *CH-Cik1 bim1Δ,* respectively) (**Figure 6B**). Further we investigated the localization of Ase1 and Bik1 in the *CH-Cik1* strain, as both are altered in the *bim1Δ* background (Figure 1B). We used Ndc80-GFP as a control since its overall levels at KT clusters did not change upon *bim1* deletion. Relative to the *bim1* deletion, expression of *CH-Cik1* decreased the amount of Ase1-GFP and increased the association of Bik1-3xGFP to the metaphase spindle, while the control Ndc80 remained unchanged (**Figure 6C, D**). Moreover, *CH-Cik1* almost fully rescued initial metaphase spindle length and partially the maximal metaphase spindle length (wild-type: 0.92±0.01 μm and 1.98±0.02 μm; *bim1Δ*: 0.67±0.01 μm and 1.48±0.03 μm; *CH-Cik1*: 0.85±0.02 μm and 1.63±0.03 μm) (**Figure 6E, F, G**). Overall, *CH-Cik1* improved metaphase spindle organization, but the cells retained a characteristic of the *bim1* deletion in the cytoplasm, displaying high levels of Bik1 on long astral microtubules (**Figure 6C**). In genetic crosses the *CH-Cik1* chimera fully rescued the lethality of the *bim1Δ ase1Δ* and *bim1Δ slk19Δ* double mutants. Remarkably, even the lethality of *bim1Δ* in cells lacking the mitotic checkpoint (*mad1Δ*) was partially rescued by the *CH-Cik1* fusion, as corresponding spores were viable, albeit small. By contrast, the *CH-Cik1* chimera was unable to rescue the lethality of *bim1Δ* mutants in combination with mutants in the nuclear migration pathway (*bik1Δ* or *jnm1Δ*), or in tension sensing and centromeric cohesion (*chl4Δ*) (**Figure 6H**). We have previously shown that the phenotype of Bim1-binding deficient Cik1 mutants can be rescued by fusing the CH-domain to this Cik1 mutant (*cik1-Δ74*). The experiments in this study extend those findings and show that a *CH-Cik1* fusion is even able to compensate efficiently for a complete lack of Bim1, at least with regards to nuclear functions.

**Figure 6.**
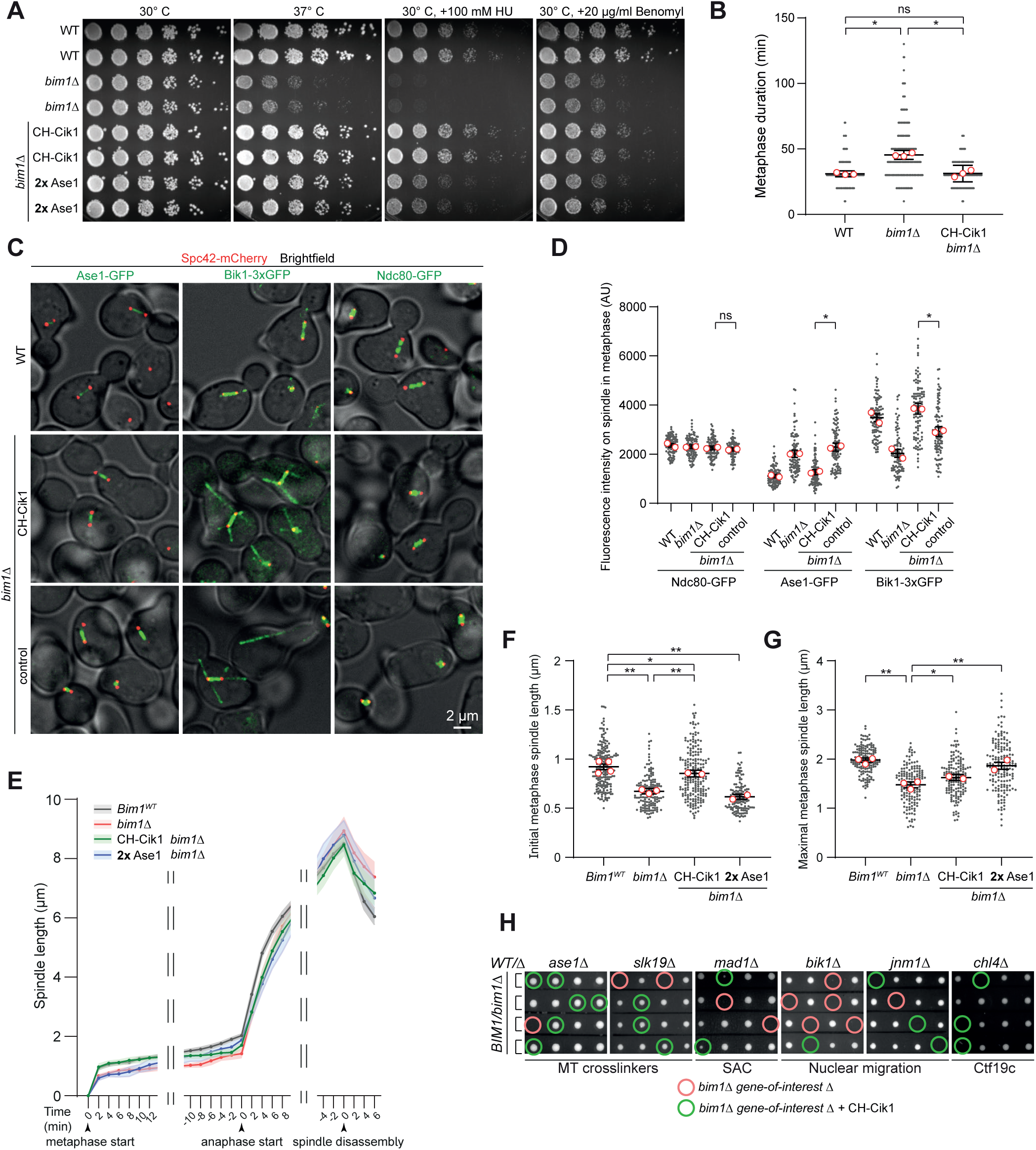
Expression of CH^Bim1^-Cik1 or increased dosage of the crosslinker Ase1/PRC1 can compensate for lack of Bim1 in the nucleus. **(A)** Serial dilution assay testing growth at 30°C, 37°C or at 30°C in the presence of 100 mM Hydroxyurea or 20 μg/ml benomyl. Plates were photographed after 2.5 days. Note the rescue of temperature-, hydroxyurea- and partial rescue of benomyl sensitivity of a *bim1Δ* mutant by expression of the CH-Cik1 fusion. **(B)** Quantification of metaphase duration in cells expressing a chimeric CH-Cik1 fusion protein or a control integration in *bim1Δ* cells. 300 cells were analyzed for each condition. Small dots correspond to individual cells, open red circles represent mean values of three biological repeats. Error bars are mean values with 95% CIs. Asterisks denote: *, P=0.0001; ns, non-significant (P=0.9893); One-way ANOVA with Tukey’s post-test was used. **(C)**. Merged images of brightfield and fluorescent microscopy of the marker proteins Ase1-GFP, Bik1-3xGFP and Ndc80-GFP in wild type cells or in *bim1Δ* cells with an integrated CH-Cik1 fusion gene versus a control. Scale bar 2 μm. **(D)** Quantification of fluorescence intensity on metaphase spindle of marker proteins Ndc80-GFP, Ase1-GFP and Bik1-3xGFP in the indicated strain backgrounds. 100 cells were analyzed for CH-Cik1 and control strains for each condition. Small dots correspond to individual cells, open red circles represent mean values of repeats. Data for wild-type and *bim1Δ* cells is shown for reference and is the same as in Figure 1B. Error bars are mean values with 95% CIs. Asterisks denote *, P<0.0001; ns, non-significant (p=0.2038); unpaired two-tailed Student’s *t* test was used. **(E)** Spindle elongation kinetics of the indicated strains. 25 cells were analyzed for CH-Cik1 and control strains, data for wild-type and *bim1Δ* cells is shown for reference and is the same as in Figure 2C. Curves show mean values with 95% CIs. Breaks in the graph correct for differences in mitotic timing. **(F)** Quantification of initial metaphase spindle length of the indicated strains. 200 cells were analyzed for CH-Cik1 and 130 cells for 2x Ase1 strains, data for wild-type and *bim1Δ* cells is shown for reference and is the same as in Figure 2E. Small dots correspond to individual cells, open red circles represent mean values of repeats. Error bars are mean values with 95% CIs. Asterisks denote: **, P<0.0001; *, P=0.0152; Welch and Brown-Forsythe ANOVA with Games and Howell post-test was used. **(G)** Quantification of maximal metaphase spindle length of the indicated strains. 75 cells were analyzed for CH-Cik1 and 100 cells for 2x Ase1 strains, data for wild-type and *bim1Δ* cells is shown for reference and is the same as in Figure 2E. Small dots correspond to individual cells, open red circles represent mean values of repeats. Error bars are mean values with 95% CIs. Asterisks denote: **, P<0.0001; *, P=0.0045; Welch and Brown-Forsythe ANOVA with Games and Howell post-test was used. **(H)** Heterozygous diploid dissection of different gene deletions in a *bim1Δ* background expressing CH-Cik1 fusion. Four spores of a tetrad are displayed horizontally, the indicated genotypes are marked by circles: Red and green circles compare growth of the respective double mutant in absence or presence of the CH-Cik1 fusion, respectively. Plates were photographed after 2 days.

### Increased dosage of Ase1 can partially substitute for the Bim1-Cik1-Kar3 complex in budding yeast mitosis

We asked to what extent elevated expression of Ase1 may rescue the growth defects of the *bim1Δ* mutant. We constructed a series of strains expressing an extra copy of Ase1-GFP under different constitutive promoters of increasing strength (p1 to p6, **Supplementary Video 3**). While high overexpression of Ase1 from the p6 promoter was lethal, moderate expression levels showed a dose-dependent rescue of temperature sensitivity of the *bim1Δ* mutant (**Supplementary Figure 6A**). Surprisingly, a single extra copy of Ase1 (*2xAse1*) expressed from its native promoter already provided a decent rescue of *bim1Δ*, albeit not as efficiently as *CH-Cik1* (**Figure 6A**, **Supplementary Figure 6A**). Exogenously expressed Ase1 displayed a similar level and kinetics of localization compared to the endogenous protein, indicating that binding sites for microtubule crosslinkers are not a limiting factor on the budding yeast spindle (**Supplementary Figure 6B**). Quantification of metaphase spindle elongation showed that elevated expression of Ase1 (*2xAse1*) was unable to rescue the increased metaphase duration and decreased initial metaphase spindle length of *bim1!1* cells, but efficiently restored maximal metaphase spindle length (0.62±0.01 μm and 1.86±0.04 μm) (**Figure 6E, F, G**). This highlights a difference in the importance of early and late microtubule crosslinking in metaphase. Early crosslinking is provided by Bim1-Cik1-Kar3 and is important for efficient chromosome bi-orientation. Late crosslinking is mediated by Ase1 and is required for development of full spindle length.

### The C-terminus of Bim1 contributes to the formation of a Bim1-Bik1-Cik1-Kar3 complex

Finally, we aimed to investigate the function of the key Bim1-cargo complexes during mitosis also from the perspective of Bim1, not relying on a *bim1* deletion mutant but on specific *bim1* mutations. To this end, we constructed mutants that interfere with different types of Bim1-dependent interactions. A double point mutation exchanging two conserved residues in the EBH domain (*bim1 Y220A E228A*) is predicted to eliminate all EBH-dependent cargo interactions, but does not affect protein dimerization. A deletion of the terminal five amino acids is predicted to prevent binding of the CAP-Gly domain of Bik1 to Bim1. The combination of both mutations is expected to simultaneously prevent both types of interaction (**Figure 7A**). First, we tested the designed Bim1 mutants *in vitro*. Recombinant proteins were well expressed and displayed identical elution profiles during size-exclusion chromatography, indicating identical dimerization status (**Supplementary Figure 7A**). As expected, the Bim1EBH protein was deficient in Cik1-Kar3 binding, but capable of binding Bik1. In contrast, Bim1^ΔC^ was able to co-elute with Cik1-Kar3, but not with Bik1. The Bim1^EBHΔC^ mutant was deficient in both types of interactions. Next, we examined the phenotypes of these *bim1* mutants *in vivo* using live cell microscopy (**Figure 7B, Supplementary Video 4**). Localization of Kar9 on bud-directed astral microtubules was lost in *bim1^EBH^* and *bim1^EBHΔC^* mutants similar to *bim1Δ*, but unaffected by *bim1^ΔC^*(**Figure 7C**). Interestingly, localization of Cik1-Kar3 on the spindle was not only dependent on direct Bim1 binding using the EBH domain but was also compromised in the *bim1^ΔC^* mutant (**Figure 7D**).

**Figure 7.**
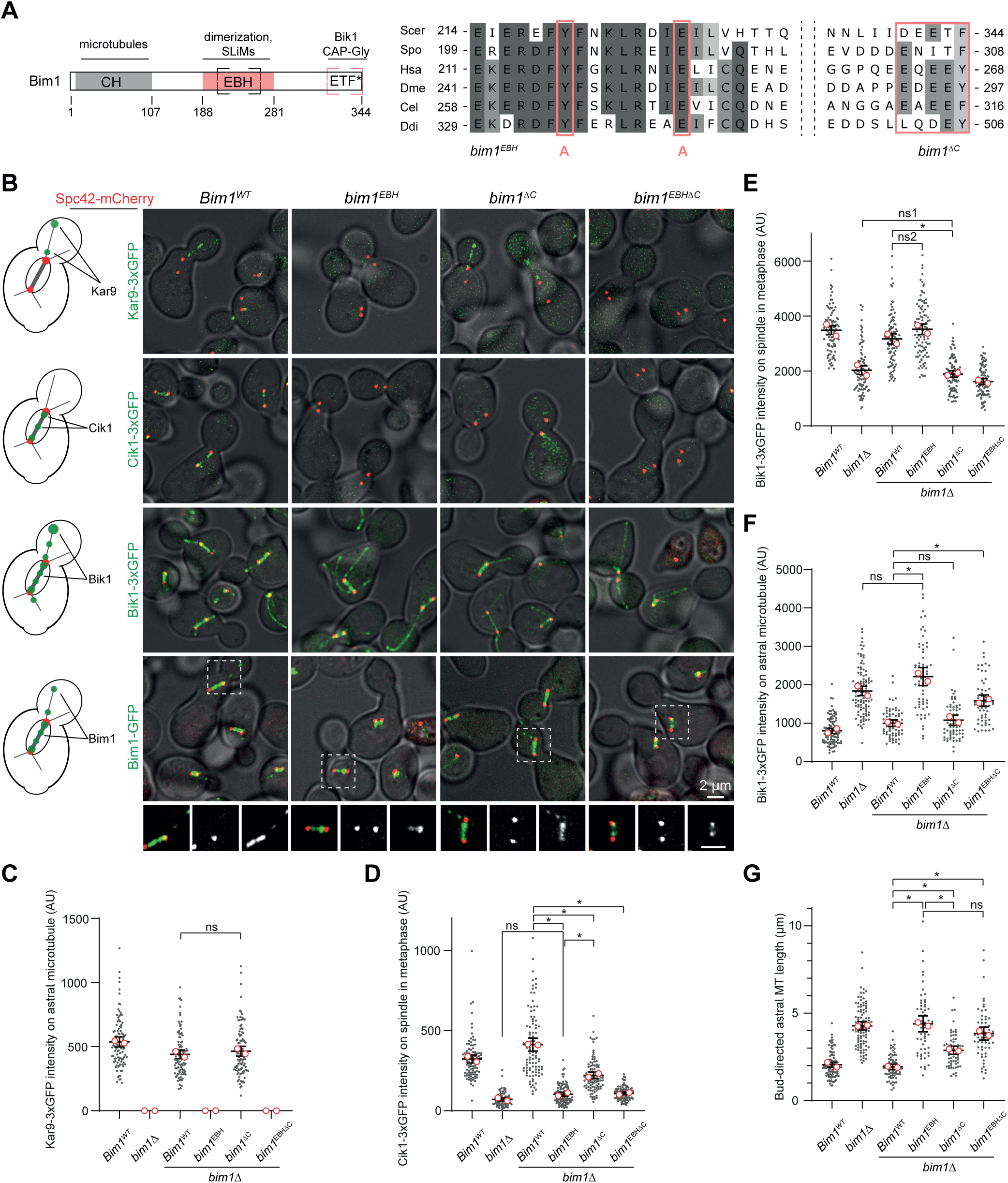
The C-terminus of Bim1 contributes to the spindle localization of Cik1. **(A)** Scheme of Bim1 and indication of point mutations introduced into the EBH domain and the deletion of C-terminal tail. Multiple sequence alignment prepared in Unipro Ugene shows part of the conserved EBH domain and C-terminus, shaded residues indicate percentage of identity. **(B)** Fluorescent micrographs of marker proteins Kar9-3xGFP, Cik1-3xGFP, Bik1-GFP in Bim1 wild-type or mutant strains. Last row depicts localization of GFP-tagged Bim1 wild-type or mutants. On the bottom, magnification of the boxed area is shown. From left to right: merged image, red channel, green channel. Strains additionally express Spc42-mCherry. Scale bar, 2 μm. **(C)** Quantification of Kar9-3xGFP intensity on astral microtubules in different *bim1* mutants. 100 cells were analyzed for each condition. Small dots correspond to individual cells, open red circles represent mean values of repeats. Data for wild-type and *bim1Δ* cells is shown for reference and is the same as in Figure 1B. No signal above background was detected in *bim1Δ, bim1^EBH^*, *bim1^EBHΔC^* cells. Error bars are mean values with 95% CIs. ns, non-significant (p=0.6361); Welch and Brown-Forsythe ANOVA with Games and Howell post-test was used. **(D)** Quantification of Cik1-3xGFP intensity of spindle microtubules in different *bim1* mutants. 100 cells were analyzed for each condition. Data for wild-type and *bim1Δ* cells is shown for reference and is the same as in Figure 1B. Small dots correspond to individual cells, open red circles represent mean values of repeats. Error bars are mean values with 95% CIs. Asterisks denote: *, P<0.0001; ns, non-significant (p=0.9986); Welch and Brown-Forsythe ANOVA with Games and Howell post-test was used. **(E)** Quantification of Bik1-3xGFP intensity on spindle microtubules in metaphase in different *bim1* mutants. 100 cells were analyzed for each condition. Data for wild-type and *bim1Δ* cells is shown for reference and is the same as in Figure 1B. Small dots correspond to individual cells, open red circles represent mean values of repeats. Error bars are mean values with 95% CIs. Asterisks denote: *, P<0.0001; ns, non-significant (ns1, p=0.7219; ns2, p=0.1074); Welch and Brown-Forsythe ANOVA with Games and Howell post-test was used. **(F)** Quantification of Bik1-3xGFP intensity on astral microtubules in different Bim1 mutants. More than 60 cells were analyzed for each condition. Data for wild-type and *bim1Δ* cells is shown for reference and is the same as in Figure 1B. Small dots correspond to individual cells, open red circles represent mean values of repeats. Error bars are mean values with 95% CIs. Asterisks denote: *, P<0.0001; ns, non-significant (p=0.9261); Welch and Brown-Forsythe ANOVA with Games and Howell post-test was used. **(G)** Quantification of length of bud-directed astral microtubules in different *bim1* mutants. More than 60 cells were analyzed for each condition. Data for wild-type and *bim1Δ* cells is shown for reference and is the same as in Figure 5F. Small dots correspond to individual cells, open red circles represent mean values of repeats. Error bars are mean values with 95% CIs. Asterisks denote: *, P<0.0001; ns, non-significant (p=0.4272); Welch and Brown-Forsythe ANOVA with Games and Howell post-test was used.

Nuclear Bik1 localization was reduced in the *bim1^ΔC^*mutant, however this did not cause an increase of cytoplasmic Bik1, arguing for the existence of at least partially independent pools of Bik1 in the nucleus and cytoplasm. In contrast to this, the *bim1^EBH^* mutant had a strong nuclear Bik1 signal, but induced the accumulation of Bik1 on the lattice of astral microtubules, similar to *bim1!1* (**Figure 7E, F**). The observed effects were not due to changes in the expression levels of the Bim1-interacting proteins (**Supplementary Figure 7E**). Next, we asked how the *bim1* mutations affect their own localization. As expected, *bim1^EBH^*displayed an exclusively nuclear localization indicating that cytoplasmic localization of Bim1 is mediated by interaction with Kar9 in mitotic cells, and in complexes with Kar9 and Bik1-Cik1-Kar3 in α-factor arrested cells. We found that all *bim1* mutants were less well recruited to the metaphase spindle compared to the wild-type protein, indicating that Bim1-interacting proteins strongly contribute to Bim1 localization (**Figure 7B, Supplementary Figure 7C, D**). To test that the observed recruitment defects of *bim1* mutants are not a result of a compromised spindle or microtubule structure, we examined their localization in a situation when GFP-tagged mutants were covered with the unlabeled wild-type allele. Indeed, in this situation, the Bim1 mutants displayed very similar localization profiles (**Supplementary Figure 7B**).

Using serial dilution assays we found that the *bim1^EBH^* mutant closely resembles the *bim1* deletion phenotype, while *bim1^ΔC^* grew indistinguishably from wild-type cells (**Figure 8A**). The phenotype of *bim1^EBH^* in growth assays was not exacerbated by additional deletion of its C-terminus. A closer analysis of the cellular phenotypes regarding metaphase duration (**Figure 8B**) confirmed these results. In an attempt to detect more subtle defects, we analyzed the spindle association dynamics of Ase1-GFP. Bim1^ΔC^ cells displayed an increased association of Ase1-GFP on nascent metaphase spindles, similar to *bim1^EBH^* and *bim1^EBHΔC^* cells. This suggests that loss of Bik1 and Bik1-Cik1-Kar3 binding contributes to the strong accumulation of Ase1 on metaphase spindles (**Figure 8C, E**). By analyzing spore viability in the absence of the SAC we confirmed that the Bim1-Bik1 interaction is required for error-free chromosome bi-orientation (**Figure 8D**). Overall, the experiments using *bim1* mutants confirm that the key functions of Bim1 are executed by complexes whose formation relies on the EBH domain alone (Bim1-Kar9), or on the EBH domain in combination with the C-terminus (Bim1-Bik1-Cik1-Kar3). The observation that *bim1*^EBH^ and *bim1*^EBHΔC^ exhibit the same phenotypes as the *bim1* deletion, indicate that without cargo-binding Bim1 cannot efficiently contribute to microtubule regulation in cells.

**Figure 8.**
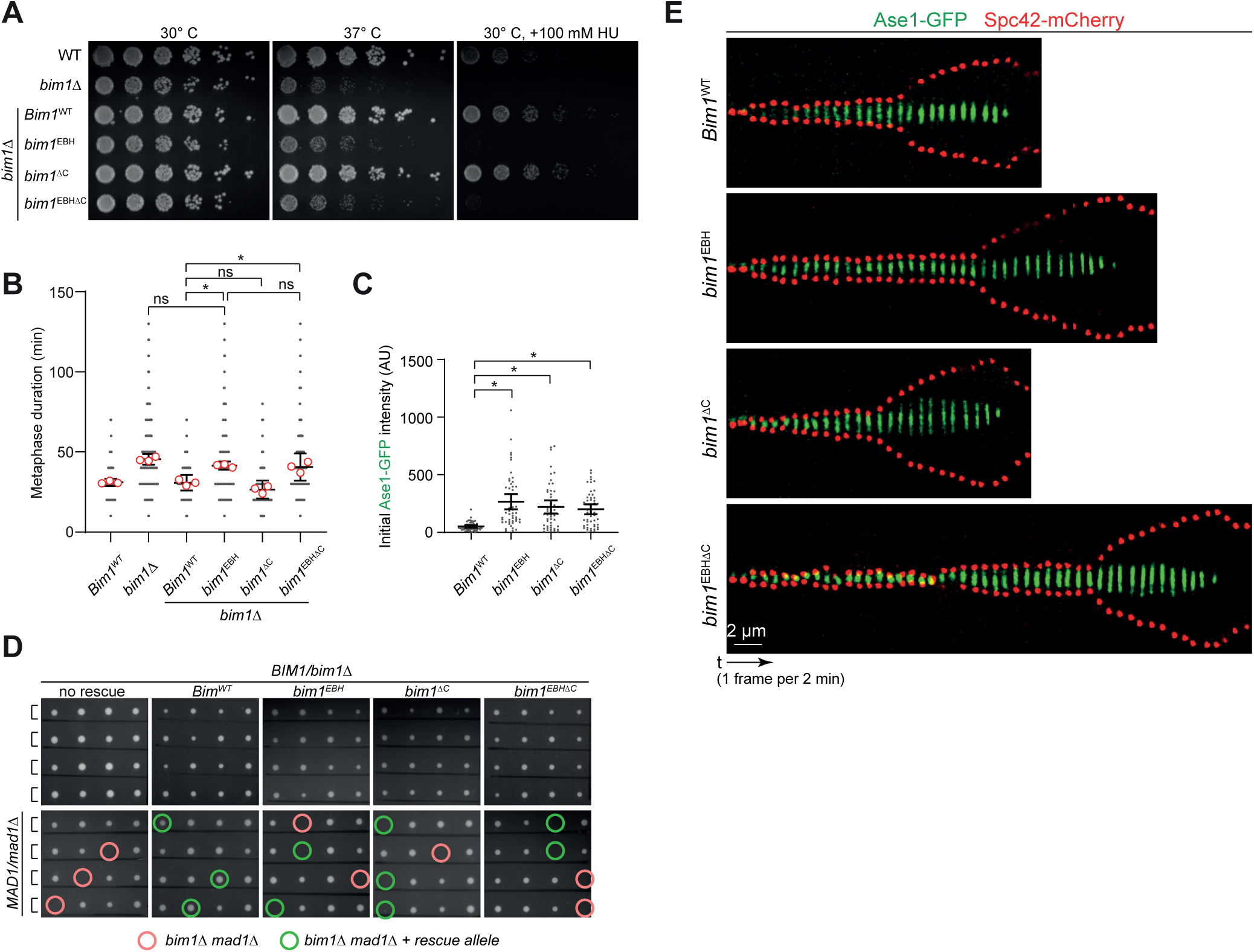
Characterization of Bim1 mutations in EBH domain and C-terminus. **(A)** Serial dilution assay of Bim1 mutants, testing growth at 30°C, 37°C and in the presence of 100 mM Hydroxyurea. Plates were photographed after 2.5 days of incubation under the indicated conditions. **(B)** Quantification of metaphase duration in different *bim1* mutants. Note that *bim1^EBH^* and *bim1^EBHΔC^* mutants closely resemble the *bim1*Δ phenotype. Data for wild-type and *bim1Δ* cells is shown for reference and is the same as in Figure 6B. 300 cells were analyzed for each condition. Small dots correspond to individual cells, open red circles represent mean values of three biological repeats. Error bars are mean values with 95% CIs. Asterisks denote: *, P<0.001; ns, non-significant; One-way ANOVA with Tukey’s post-test was used. **(C)** Quantification of initial Ase1-GFP intensity on metaphase spindles in different *bim1* mutants. More than 50 cells were analyzed for each condition. Small dots correspond to individual cells. Error bars are mean values with 95% CIs. Asterisks denote: *, P<0.0001; Welch and Brown-Forsythe ANOVA with Games and Howell post-test was used. **(D)** Heterozygous diploid dissection of *bim1* mutants in *bim1Δ* background (top row), or in a *bim1Δ mad1Δ* background (bottom). Four spores of a tetrad are displayed horizontally; the indicated genotypes are marked by circles. Note lethality of *bim1* cargo-binding mutants in the *mad1Δ* background. **(E)** Kymographs displaying Ase1-GFP association dynamics to metaphase spindles in wild-type or different *bim1* mutant cells. Timelapse 2 min, scale bar 2 μm.

## Discussion

In this study we have comprehensively analyzed the role of the sole EB protein Bim1 in budding yeast mitosis using a combination of quantitative live cell imaging and genetic analysis. Our experiments allow us to unify previous observations into a more comprehensive picture of Bim1-dependent and -independent processes during mitosis and they help to explain how cells organize a microtubule cytoskeleton in the absence of an EB protein.

### Defining a minimal set of Bim1 cargo complexes required for efficient mitosis in budding yeast

Successful chromosome segregation in the closed mitosis of budding yeast requires two microtubule-based processes: 1. Assembly of a bipolar intranuclear spindle on which sister chromatids bi-orient and are then separated as the spindle elongates in anaphase. 2. Positioning of the yeast nucleus along the bud-neck axis such that the elongating spindle distributes one set of sister chromatids into mother and daughter cell. Our analysis indicates that Bim1 contributes to both of these processes as part of two key protein complexes (**Figure 9A**): Bim1-Kar9-Myo2 in the cytoplasm and Bim1-Bik1-Cik1-Kar3 in the nucleus. These complexes differ from other Bim1 interaction partners in several features: Kar9 and Cik1-Kar3 both strictly require Bim1 for microtubule localization as judged by live cell microscopy. This behavior is characteristic, as none of the other 20 tested GFP-fusions were as severely affected by the *bim1* deletion. Bim1 and Kar9 mutually depend on each other for localization to cytoplasmic microtubules. In the nucleus, Bim1 is able to localize to the spindle in the absence of Cik1 or Kar3, but the organization of the resulting spindles is as severely impaired as in the *bim1Δ* strains. It is noteworthy that formation of both of these Bim1-cargo complexes does not depend on canonical SxIP motifs, but primarily relies on non-SxIP motifs (LxxPTPh in the case of Kar9 and KLTF-ELN in the case of Cik1-Kar3) in combination with SxIP motifs. Biochemical experiments suggest that the resulting binding mode allows the formation of particularly stable Bim1-cargo complexes. The notion that Bim1 executes its cellular functions as part of these stable complexes is supported by the observation that the *bim1^EBH^* mutant essentially displays the same phenotypes as the *bim1* deletion in our analysis. Thus, microtubule plus-end binding by Bim1 itself contributes little to overall microtubule regulation in cells and instead Bim1 executes its functions as part of stable cargo complexes.

**Figure 9.**
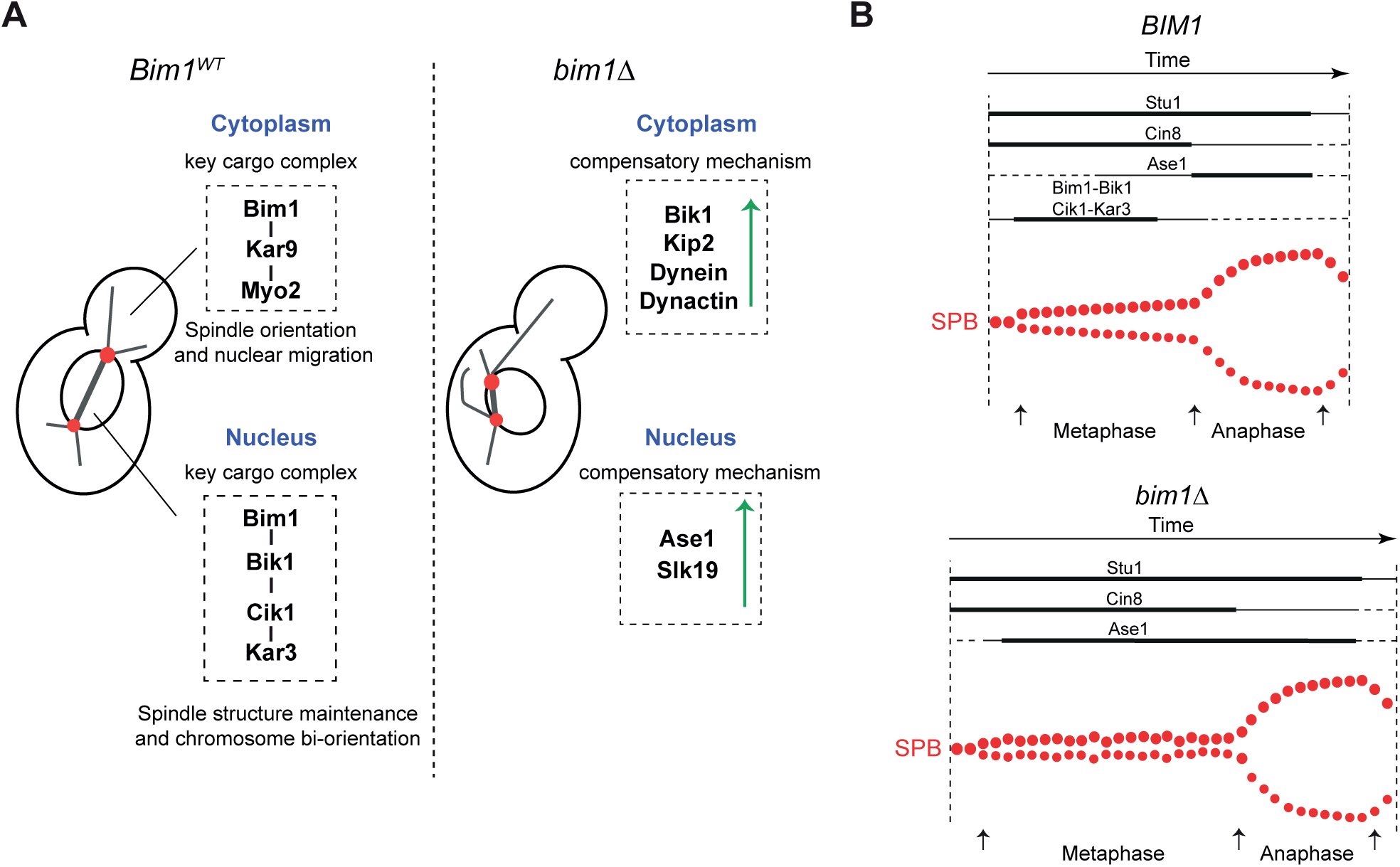
Model for functions of key Bim1-cargo complexes. **A:** Key Bim1 cargo complexes in cytoplasm and nucleus and parallel pathways required in their absence. In the *bim1Δ* strain, upregulation of the Dynein-Dynactin pathway is required for nuclear migration. In the nucleus, increased association of Ase1 or Slk19 is required for metaphase spindle assembly in the absence of Bim1. **B:** Schematic representation of spindle assembly and timing of association of microtubule-associated proteins in wild-type and *bim1* deletion cells. See discussion for details.

Our study highlights the notion that a lack of EB has similar overall consequences on microtubule organization in different systems. Similar to our findings in budding yeast, acute photo-inactivation of EB in human cells leads to a shortening of the metaphase spindle, while anaphase phenotypes are comparatively minor (Dema et al., 2022). Similar effects on spindle length have been reported upon EB RNAi in Drosophila cells (Goshima et al., 2007; Li et al., 2011). An increase in astral microtubule length, similar to the effects of a *bim1!1*, has also been reported as a consequence of EB inactivation in human cells (Dema et al., 2022). In this case a lack of EB-Kif18B complexes is responsible for the impaired astral microtubule length control. Thus, even though overall consequences of EB loss-of-function maybe similar, they can involve different EB-cargo complexes in different systems.

### Rescuing Bim1-dependent spindle phenotypes with a CH-Cik1 fusion protein

A surprising finding from our analysis is that despite many potential binding partners in yeast, the spindle-related phenotypes of a *bim1* deletion can be efficiently rescued by providing a single Bim1 cargo – the Kinesin-14 motor Kar3 in the form of a chimeric *CH-Cik1* fusion protein. In particular, our analysis indicates that the *CH-Cik1* fusion rescues the increased metaphase duration of a *bim1* deletion strain, restores wild-type initial metaphase spindle length, decreases the amount of Ase1 on the metaphase spindle (indicative of normal spindle organization) and re-establishes the organization of kinetochore clusters. There are, however, two indications that the *CH-Cik1 bim1Δ* strain does not fully behave like a wild-type. As it lacks Bim1-Kar9 complexes in the cytoplasm, it has a spindle positioning defect. This may partly explain the incomplete rescue of the *bim1Δ mad1Δ* strain. Besides, the double mutant *bim1Δ chl4Δ* remains inviable even upon expression of *CH-Cik1*. Perhaps the chimera is very effective in providing efficient spindle organization, indicated by its ability to rescue *bim1Δ ase1Δ* or *bim1Δ slk19Δ* double mutants, but cannot fully compensate for Bim1’s roles in outer kinetochore function or tension-dependent sister chromatid bi-orientation. Future experiments will have to relate these observations to the precise molecular functions of Bim1-Cik1-Kar3.

### Differential contributions of Bim1 and Bik1 to parallel spindle orientation pathways

Spindle positioning in budding yeast is achieved via two pathways, one relying on the protein Kar9 which interacts with the actin-based motor Myo2 (Hwang et al., 2003), the other involving the microtubule-based motor cytoplasmic Dynein (Li et al., 1993). For nuclear migration to occur efficiently, the Dynein-Dynactin complex must be enriched at the plus-ends of cytoplasmic microtubules, contrary to its intrinsic minus-end directed motility (Reck-Peterson et al., 2006). While *in vitro* reconstitution experiments have suggested that Bim1 is required to fully reconstitute the Kip2-dependent loading of the Dynein-Dynactin complex to microtubule-plus ends *in vitro* (Roberts et al., 2014), our experiments indicate that it may contribute relatively little to this pathway in cells. If this was the case, then the nuclear migration defect of *bim1* deletion cells should be much more pronounced than observed and stronger than in *kar9*!1 cells. The dominating effect of the *bim1* deletion with regards to the Dynein-Dynactin pathway, however, appears to be the increase in Bik1 on long cytoplasmic microtubules. Bik1 foci move towards the plus end with a velocity of 6-12 μm/min, corresponding to the speed of Kip2 molecules (**Figure 5G**). We speculate that in the *bim1Δ* strain Bik1 increases the processivity of Kip2, which in turn further enriches Kip2 at plus-ends, promotes the polymerization of long cytoplasmic microtubules and helps to load high levels of Dynein-Dynactin onto microtubule-plus ends. These long microtubules can interact with the bud cortex and initiate pulling events to move the nucleus (Omer et al., 2018). Bik1 binding may additionally have a more direct role in stabilizing these microtubules. We currently do not fully understand why simply re-localizing Bik1 with the *Bik1-NES* construct does not appear to lead to the formation of long microtubules in wild-type cells. It is possible that the presence of Bim1 in wild-type cells supports a counteracting activity which helps to keep cytoplasmic microtubules short.

While Bik1 has a key role in the cytoplasm that can be executed independently of Bim1, its contributions to nuclear spindle function in budding yeast are comparatively less significant and subsequent to (or downstream of) Bim1. This is supported by the observation that *bim1^11C^* cells do not display an increased metaphase duration. Lack of the Bim1 C-terminus also does not substantially aggravate the growth defects of the *bim1^EBH^* mutant, and chromosome bi-orientation and spindle elongation can occur, albeit with a delay, in the simultaneous absence of Bim1 and Bik1, as long as microtubule crosslinkers such as Ase1 are still present (**Figure 4A**). It is likely that Bik1 contributes to the nuclear functions of Bim1 mainly as an additional subunit of the Bim1-Cik1-Kar3 complex. This is indicated by the decrease of Cik1 spindle association in the *bim1^11C^*mutant and the ability of Bik1 to bind to Bim1-Cik1-Kar3 complexes *in vitro*. Future experiments will have to analyze how inclusion of Bik1 may modulate the activities of the Bim1-Cik1-Kar3 complex.

### Strategies for balancing microtubule plus-end interaction networks in cells

Our experiments reveal more general strategies that yeast cells employ to compensate for the absence of an EB protein and re-organize plus-end interaction networks. The lack of Bim1 leads to a striking intracellular redistribution of Bik1. Imaging the Bim1^WT^-GFP construct indicates that Bim1 appears to be primarily nuclear and spindle-associated. The cytoplasmic localization is restricted to individual foci and depends on Kar9. The increased association of Bik1 with cytoplasmic microtubules is required for nuclear migration in the absence of Bim1. This highlights the notion that nucleo-cytoplasmic distribution relies both on nuclear import and export signals in individual proteins, but also on the distribution of binding partners that can shift the equilibrium between protein pools in different compartments (Schweiggert et al., 2016).

A second consequence of the lack of Bim1 is an altered timing and overall increased spindle association of other MAPs, most notably of the PRC1 homolog Ase1. Ase1/PRC1 proteins are known to preferentially crosslink antiparallel microtubule bundles both *in vitro* and *in vivo* (Subramanian et al., 2010), therefore we speculate that *bim1!1* mutants may have more extended antiparallel overlap zones during early spindle assembly than wild type cells. In budding yeast 32 out of ∼40 nuclear microtubules are connected to kinetochores and have a short length of ∼340 nm (Winey et al., 1995). One possibility is that *bim1Δ* cells have a defect in the control of kinetochore microtubule length and more microtubules cross the spindle equator to the opposite kinetochore cluster (**Supplementary Figure 2F**). A similar activity has been proposed for the kinesin-5 Cin8 (Gardner et al., 2008). On monopolar spindles that would create a higher fraction of intersecting microtubules and therefore larger antiparallel overlapping zones. This could also explain the problems these cells have in establishing tension across sister kinetochores, evidenced by their high levels of Sgo1 recruitment. Our observations are consistent with the idea that ordered recruitment of different microtubule crosslinkers to the spindle reflects their functional contributions to distinct processes (**Figure 9B**). We propose that Bim1-Bik1 in a complex with its cargo Cik1-Kar3 works after bi-polar spindle formation but before late metaphase and is critical for sister chromatid bi-orientation. Ase1, on the other hand, is recruited later, stabilizes the antiparallel overlap zone of interpolar microtubules and is required for maturation of the metaphase spindle to its full pre-anaphase length. Our data also indicate that the decrease in metaphase spindle size in *bim1Δ* cells is likely not the key reason for a delay in metaphase progression and chromosome bi-orientation. Indeed, even nascent bi-polar spindles have a length of ∼0.9 μm that in principle should allow them to establish tension between sister chromatids (kinetochore microtubule length is ∼340 nm). The *ase1* depletion affects spindle length as severely as *bim1!1*, but has little effect on Sgo1 level and metaphase progression because chromosome bi-orientation is achieved largely before Ase1 recruitment on the spindle in late metaphase. Ase1 can partially substitute for nuclear Bim1 functions, but *2xAse1 bim1!1* cells still have an increased metaphase duration, indicating that they are not as effective in achieving bi-orientation as wild-type cells. Collectively, our observations reinforce the notion that multiple redundant mechanisms ensure the robustness of spindle assembly and positioning. While the stereotypic construction of the yeast spindle allows a detailed quantification of spindle parameters, a current experimental limitation is that its small size prevents a more precise definition of phenotypes by light microscopy. In the future cryo-EM tomography in cells coupled with acute genetic perturbations and biochemically defined mutants promises to reveal further insights into the mechanisms by which Bim1-cargo complexes contribute to specific aspects of spindle assembly and function.

## Materials and Methods

### Yeast genetics

All strains presented in this study are part of the NKY collection. Most strains were derived by modification of the diploid strain NKY982-2 (MATa/MATα ADE2/ADE2 lys2-801/lys2-801 leu2-3,112/leu2-3,112 Spc42-mCherry::HIS3/SPC42 his3Δ200/his3Δ200 ura3::pGAL-osTIR::URA3/ura3::pGAL-osTIR::URA3 bim1Δ::hphNT1/BIM1) which is a progeny of DDY1102 in the S288c background. One clone of one repeat of every strain used in this study is listed in Supplementary Table 3. Genetic modifications were generated by standard methods (Janke et al., 2004; Longtine et al., 1998) and introduced into yeast using LiAc transformation (Schiestl and Gietz, 1989).

### Growth conditions

Yeasts were maintained in YEPD medium at 30°C unless otherwise stated. For protein depletion experiments yeasts were grown in YEP medium/plates with raffinose and galactose (2% + 2%) as a sugar source at 30°C. The following antibiotics were used for selection of desired modifications: G418, Nourseothricin (ClonNAT), Hygromycin B. Auxotrophy selection markers were Leucine, Uracil, Histidine.

### Strain construction

After modification of a diploid strain for an antibiotic selection, yeasts were first incubated on a YEPD plate and allowed to grow overnight. The next day the plate was replicated on desired selection media and let grown for 2 days until single colonies of ∼3 mm in size were visible. Typically plates had 50-100 colonies, 8-16 of them were re-streaked again on a selection plate to confirm resistance. Clones were checked for successful integration by PCRs using gene-specific and resistance cassette primers. For integrations of fluorescent protein tags the presence of signal was additionally checked at the microscope. Clones were assayed for fitness compared to a parental strain using spot assays. Further two clones were processed for sporulation and tetrad dissection. 12 tetrads from every clone were genotyped by spotting on testing plates. For chosen spores tetrads of origin were analyzed for 2-2 segregation of markers by PCR to avoid aneuploidies. Gene deletions were confirmed by PCRs for the absence of wild-type gene and presence of selection marker at correct genomic loci. Functionality of gene tagging was evaluated by comparing growth in wild-type and *bim1Δ* genetic backgrounds to corresponding untagged versions. For the experiment spores were directly taken from dissection plates and were allowed to grow in YEPD overnight. Different clones that originated from the same diploid transformation were considered as technical repeats. Transformations performed independently at different days were counted as biological repeats.

### Conditional protein depletion by the AID system

The Ubiquitin ligase osTIR1 was integrated into the URA3 locus under the control of a galactose inducible promoter (Nishimura et al., 2009). A synthetic analog of auxin NAA was used at a concentration of 1 mM to induce depletion. Below the preparation of NAA for the experiment is briefly described. 1 M NAA stock was derived by dissolving NAA in DMSO and further sterile-filtered using 0.2 μm pore size filter. Aliquots were stored protected from light at -20° C until usage. Before the experiment an aliquot was thawed and diluted 10-fold in DMSO. To neutralize pH equal volume of 0.1 M NaOH was added to the solution.

### Western blot sample preparation

Yeast cell extracts were prepared using alkaline treatment as described before (Kushnirov, 2000). Briefly, an equivalent of 2 OD_600_ of exponentially growing cells was harvested, washed with H_2_O and resuspended in 200 µl of 0.1 M NaOH. After 5 minutes incubation at room temperature cells were centrifuged again. The pellet was resuspended in 50 µl 1x SDS sample buffer and boiled for 5 minutes. 8-10 µl of lysate was loaded per gel lane.

### Used antibodies and visualization

*α*-GFP (*Roche*, from mouse) 1:1000; *α*-myc (*Covance*, 9E10 from mouse) 1:1000; *α*-tubulin-HRP 1:1000 (*Santa Cruz,* from rat); HRP-coupled *α*-mouse (*GE Healthcare*) 1:10000. All antibodies were diluted in 5% (w/v) milk in TBST. For visualization Amersham™ ECL Prime Western Blotting Detection Reagent (*GE Healthcare*) was used. Images were acquired by Amersham™ Imager 600 (*GE Healthcare*).

### Plasmid construction

All plasmid generated and used are listed in Table S3.

### Recombinant protein work and biochemistry

Protein expression, purification, size-exclusion chromatography, quantitative pulldowns were performed as described (Kornakov et al. 2020) except following modifications. Quantitative pulldowns were made in buffer containing 300 mM NaCl to better reflect physiological ion strength in yeast (van Eunen et al., 2010). Imidazole concentration was reduced from 10 to 5 mM to prevent leakage of bound proteins from TALON beads.

### Serial dilution spot assays

Yeast strains were grown overnight (16-18 h) in YEPD media. Aliquots were spun down, washed 2 times with sugar-free minimal media and resuspended in it. 1: 4 serial dilutions from a starting OD 0.2 were prepared in a sterile 96-well plate using minimal media as a solvent. Yeasts were transferred to the desired plates using 48-pin replicator VP407AH (V&P Scientific, Inc). Plates were incubated at indicated conditions for 2 – 2.5 days until colonies of ∼2 mm size were visible.

### Protein engineering and genetic constructs

#### 1. Construction of a functional GFP-tagged version of Bim1

A small library of N-, C- and internally GFP-tagged versions of Bim1 was prepared. GFP was inserted using (GS)_n_ and (GA)_n_ flexible linkers between the Bim1 coding sequence and GFP. Constructs-encoding plasmids were integrated into the LEU2 locus in the heterozygous *BIM1/bim1Δ MAD1/mad1Δ* (NKY1048-1) diploid strain. Functionality was addressed by the ability of GFP-tagged constructs to rescue temperature-, hydroxyurea- and benomyl-hypersensitivities of bim1 deletion and by viability in the absence of a functional SAC (*mad1Δ*). Tested constructs displayed a variety of phenotypes and associated defects. The only fully functional version was Bim1^WT^-GFP that contained GFP inserted close to the extreme C-terminus of Bim1 10 amino acids before the stop codon. The sequence of the insertion point is the following: <u>EGEVGVS</u>-(GA)_5_-GFP-(GGGS)_2_-<u>NNLIIDEETF</u>*; Bim1 sequences are underlined.

#### 2. Artificial relocalization of Bik1

To target Bik1 to the nucleus or the cytoplasm it was modified with NLS-GFP or NES-GFP at its native locus. Constructs used for homologous recombination were prepared using overlapping-extension PCRs. The SV40 NLS (PKKKRKV) or canonical NES (LAEKLAGLDIN) sequences were inserted directly following the Bik1 coding sequence (-NQQFF*) without a linker. Bik1-NES had an exclusively cytoplasmic localization; the Bik1 signal on the mitotic spindle was reduced to background levels. Bik1-NLS had a predominantly nuclear localization. However a weak signal corresponding to ends of short cytoplasmic microtubules was still detectable, indicating an existence of natural determinants of cytoplasmic localization of Bik1.

#### 3. Artificial plus-end targeting of Kinesin-14 (CH-Cik1)

Kinesin-14 was artificially targeted to the microtubule plus-ends by fusing the CH-domain of Bim1 to the N-terminus of the kinesin-associated protein Cik1. Briefly, amino acids 1 - 133 of Bim1 were fused to the full coding sequence of Cik1 using a (GGGS)_2_ linker. The construct contained the 5’UTR of Cik1. The difference to the previously used CH-Cik1^Δ74^ version is that CH-Cik1^FL^ retains a natural NLS and degron sequences located within the Cik1 tail. In our hands it behaves slightly better than CH-Cik1^Δ74^ and allows us to release cells from α-factor arrest more efficiently. The CH-domain of Bim1 comprises the N-terminal 133 amino acids.

#### 4. Constitutive promoter library

A small library of 6 constitutive promoters of increasing strength was constructed from the weakest p1 to the strongest p6 (p1=pREV1 < p2=pPSP2 < p3=pRNR2 < p4=pRET2 < p5=pRPL18B < p6=pCCW12). In our hands p3 provided stronger expression of Ase1-GFP than p4. All promoters were 700 bp in length, besides pPSP2 that was 400 bp.

### Live cell fluorescence microscopy

#### 1. Microscopy setup

Live cell microscopy was performed using the DeltaVison Elite widefield microscope system (*GE Healthcare*) at 30°C. A 100x objective with 1.4 NA (*Olympus*) and immersion oil with a refractive index of n = 1.518 were used. Images were typically acquired by OAI (optical axis integration) scans over 3 μm width and captured with an sCMOS camera. Images were taken every 10 min, for higher time resolution experiments every 2 min. Cells were immobilized on Concanavalin A coated 8 and 4 Well ibidi Glass Bottom μ-Slide 1.5H. All images were deconvolved using SoftWoRx 7 (options: enhanced ratio, aggressive), image analysis was performed using ImageJ/Fiji software.

#### 2. Quantification of fluorescence intensity

Fluorescence intensity was quantified as a background-corrected area under the curve derived by line scan; line scan was of 10 pixel width across the spindle or a bud-directed astral microtubule.

#### 3. Live cell imaging preparations

Cells were grown overnight (16-18 h) in YEPD media at 30°C; cultures typically achieve 24-26 OD_600_. In the morning cultures were diluted 1 to 100 in fresh media and cells were allowed to achieve exponential growth for 2-3 h at 30°C. For protein depletion experiments cells were grown in YEPRG (2% of raffinose and 2% of galactose w/v). Then, 0.01 mg/ml of α-factor was added for 2h to arrest cells in a G1-like state. Synchronized cells were spun down, α-factor was washed away with pre-warmed media 2 times for YEPD cultures and 3 times for YEPRG cultures. For protein depletion the last wash contained 1 mM of NAA. After the last wash cells were resuspended in doTRP minimal media with a sugar source as desired.

## Statistical analysis

Statistical tests and graphs were prepared using Graphpad Prism 8 and 9. Small points on graphs show values of individual cells, large open circles represent means of biological or technical replicates, as indicated. A 95% Confidence Interval (CI) is usually shown. In the text mean values ±SEM are mentioned. Student’s t-test was used to compare continuous data from two strains. For multiple comparisons Welch and Brown-Forsythe ANOVA with Games and Howell post-test was used to compare pooled data; One-way ANOVA with Tukey’s post-test was used to compare data with biological repeats, as indicated.

## Acknowledgements

The authors acknowledge Alexander Dudziak, Christian Cozma and Jennifer Harris for critical reading of the manuscript and all members of the Westermann lab for discussions. Microscopy experiments were carried out with support from the Imaging Center Campus Essen Core Facility (ICCE). The DeltaVision Elite high resolution microscope was obtained through Deutsche Forschungsgemeinschaft funding (Major Research Instrumentation Programme as per Art. 91b GG, INST 20876/275-1). S.W acknowledges funding by the Deutsche Forschungsgemeinschaft (DFG, German Research Foundation)-SFB1430-Project-ID 424228829.

## Author contributions

N.K. conducted all genetic, biochemical and live cell imaging experiments. N.K and S.W. wrote the manuscript, S.W. supervised the study and acquired funding.

## Description of online supplemental material

**Table S1** contains supplementary data for microscopy-based screen of putative Bim1-binding proteins in *S.cerevisiae*. Data was prepared using www.kegg.jp; (Kanehisa and Goto, 2000). Structured and unstructured protein regions were judged by AlphaFold2 predictions (Jumper et al., 2021).

**Table S2** contains source data for **Figure 3H**. Data was prepared using TheCellMap.org; (Costanzo et al., 2016) (van Leeuwen et al., 2020).

**Table S3** contains a list of budding yeast strains and plasmids used in this study.

**Video 1** shows localization dynamics of several microtubule-associated proteins (Bim1^WT^-GFP, Bik1-3xGFP, Stu1-GFP, Ase1-GFP) in wild-type and *bim1Δ* cells over progression through mitosis. Spindle pole was visualised with Spc42-mCherry. Timelapse 2 min, scale bar 2 μm. Video is shown at 5 frames/s.

**Video 2** shows localization dynamics of several kinetochore-associated proteins (Stu2-GFP, Ndc80-GFP, Sgo1-GFP) and cytoplasmic kinesin Kip2-3xGFP in wild-type and *bim1Δ* cells over progression through mitosis. Spindle pole was visualised with Spc42-mCherry. Timelapse 2 min, scale bar 2 μm. Video is shown at 5 frames/s.

**Video 3** shows a partial rescue of *bim1Δ* with elevated expression of Ase1-GFP from promoters of increasing strength. Spindle pole was visualised with Spc42-mCherry. Timelapse 2 min, scale bar 2 μm. Video is shown 5 frames/s.

**Video 4** shows localization dynamics of several microtubule-associated proteins in *bim1*-mutant strains. Spindle pole was visualised with Spc42-mCherry. Timelapse 2 min, scale bar 2 μm. Video is shown 5 frames/s.

---

**Supplementary Figure S1.**
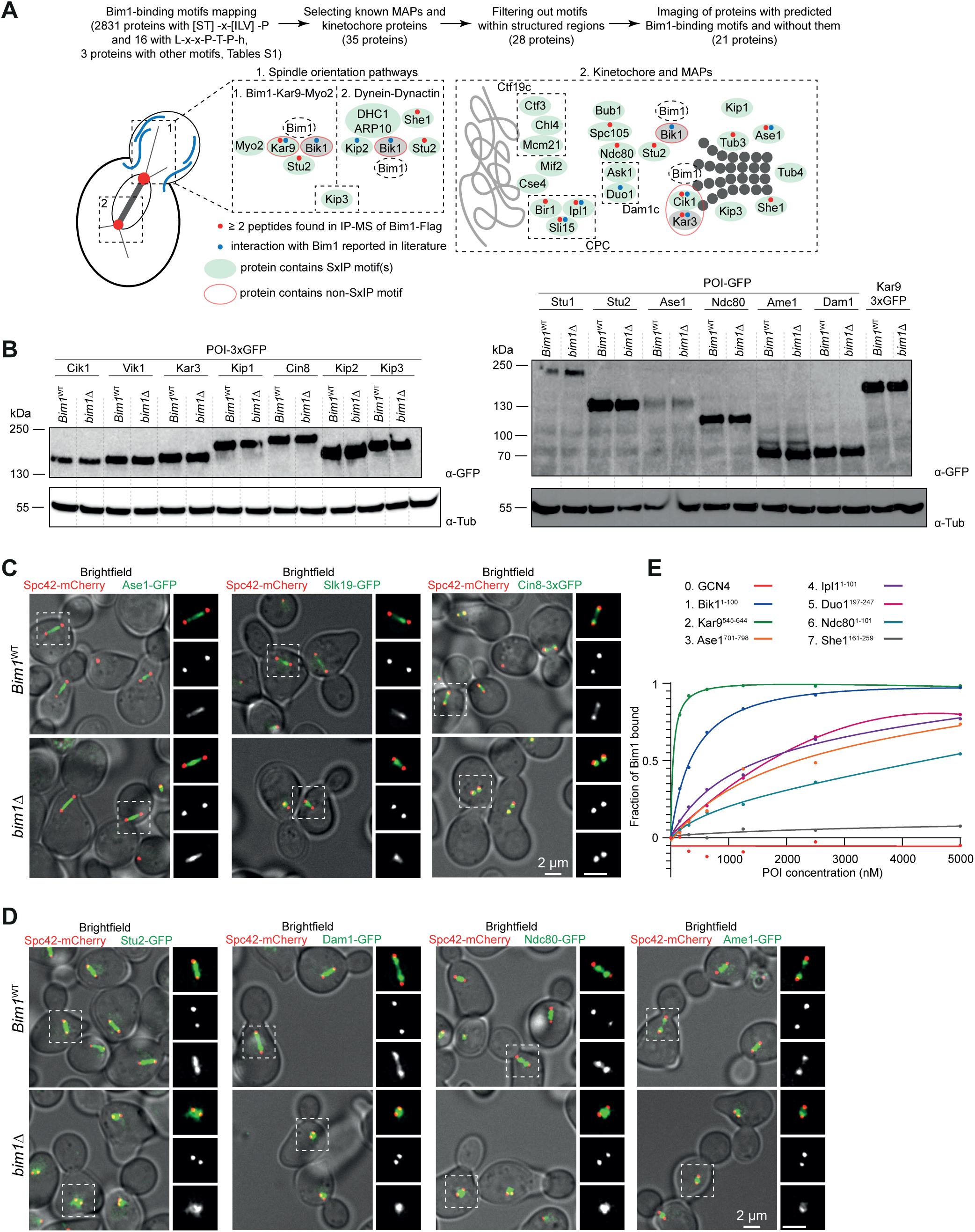
Additional characterization of GFP-Fusion proteins. **(A)** Scheme describing the stepwise selection of potential Bim1 interaction partners for analysis by live cell microscopy. **(B)** Western Blot analysis of expression of 3xGFP or GFP-tagged candidate protein in wild-type or *bim1Δ* background. α-Tubulin blots are shown as loading control. **(C)** Additional examples for the effects of *bim1*Δ on the localization of microtubule crosslinkers in the nucleus. Left side: Overlay of fluorescence microscopy with bright field image. On the right side of every image magnification of the boxed area is shown. From top to bottom: merged image, red channel, green channel. Scale bar, 2 μm. **(D)** Additional examples for the effects of *bim1Δ* on the localization of different kinetochore-associated proteins. Left side: Overlay of fluorescence microscopy with bright field image. On the right side of every image magnification of the boxed area is shown. From top to bottom: merged image, red channel, green channel. Scale bar, 2 μm. **(E)** Binding curves displaying the affinity of different GCN4 reporter constructs for Bim1 in pull-down assays *in vitro*. Binding is well approximated with the one-site total binding model. Curve fit assumes one-site total binding, which includes specific binding and a linear nonspecific binding component: Y = B_max_ * X/(*K*_d_ + X) + NS * X + Background, where Y is total binding, X is ligand, *K*_d_ is dissociation constant, NS is the slope of unspecific binding, and B_max_ is the maximum specific binding. Curves are averages of three biological repeats. Error bars are not shown for clarity.

**Supplementary Figure S2.**
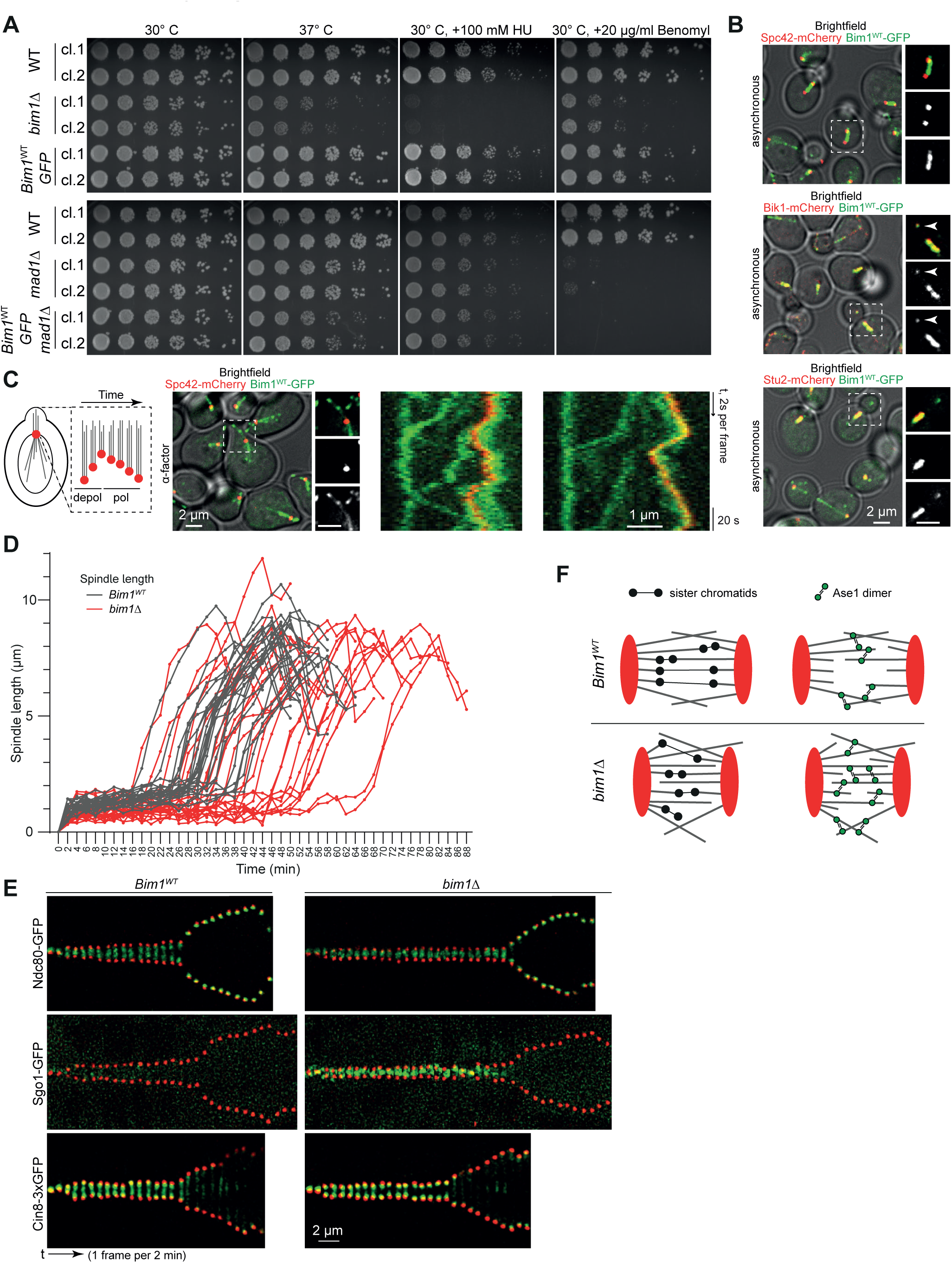
Additional characterization of the functionality of the Bim1^WT^-GFP fusion protein. **(A)** Serial dilution assays comparing growth characteristics of Bim1^WT^-GFP to wild-type or *bim1Δ* strains. cl., denotes clones 1 and 2, respectively. Plates were incubated under the indicated conditions and photographed after 2.5 days. Note that Bim1^WT^-GFP does not display temperature-, hydroxyurea-or benomyl-sensitivity and displays only mildly compromised growth at elevated temperature in combination with a checkpoint deletion (*mad1Δ*). **(B)** Fluorescence micrographs analyzing co-localization between Bim1^WT^-GFP and different markers Spc42-mCherry, Bik1-mCherry and Stu2-mCherry. Left side: Overlay of fluorescence microscopy with bright field image. On the right side of every image magnification of the boxed area is shown. From top to bottom: merged image, red channel, green channel. Arrowheads indicate co-localization of Bim1^WT^-GFP and Bik1-Cherry on plus-ends of cytoplasmic microtubules. Scale bar, 2 μm. **(C)** Analysis of Bim1^WT^-GFP dynamics on cytoplasmic microtubule bundles in α-factor arrested cells. Cycles of bundle shrinkage and growth cause pulling and pushing of the nucleus with associated SPB toward and away from the shmoo tip. Overlay of fluorescence microscopy with bright field image. On the right side magnification of the boxed area is shown. From top to bottom: merged image, red channel, green channel. Scale bar, 2 μm. Kymograph analysis of Bim1^WT^-GFP and Spc42-mCherry on shmoo tip directed microtubule bundle. Scale bar, 1 μm; time-lapse 2 s/frame. Note that Bim1^WT^-GFP tracks both polymerizing and depolymerizing ends of the microtubule bundle and individual microtubules growing along the bundle. **(D)** Example of spindle elongation analysis in wild-type and *bim1Δ* strains uncorrected for different durations of metaphase. 25 curves of wild-type spindles (grey) and *bim1Δ* spindles (red) were overlaid on the same graph. Note that *bim1Δ* metaphase spindles tend to be shorter in length and cells typically spend more time in metaphase. The analysis relates to Figure 2C. **(E)** Kymograph analysis of additional marker proteins Ndc80-GFP, Sgo1-GFP and Cin8-3xGFP in wild-type and *bim1Δ* cells. Note high levels of Sgo1-GFP on metaphase spindle in the *bim1Δ* mutant. Scale bar 2 μm, 1 frame/2min. **(F)** Schematic representation of spindle organization in wild-type and *bim1Δ* cells.

**Supplementary Figure S5:**
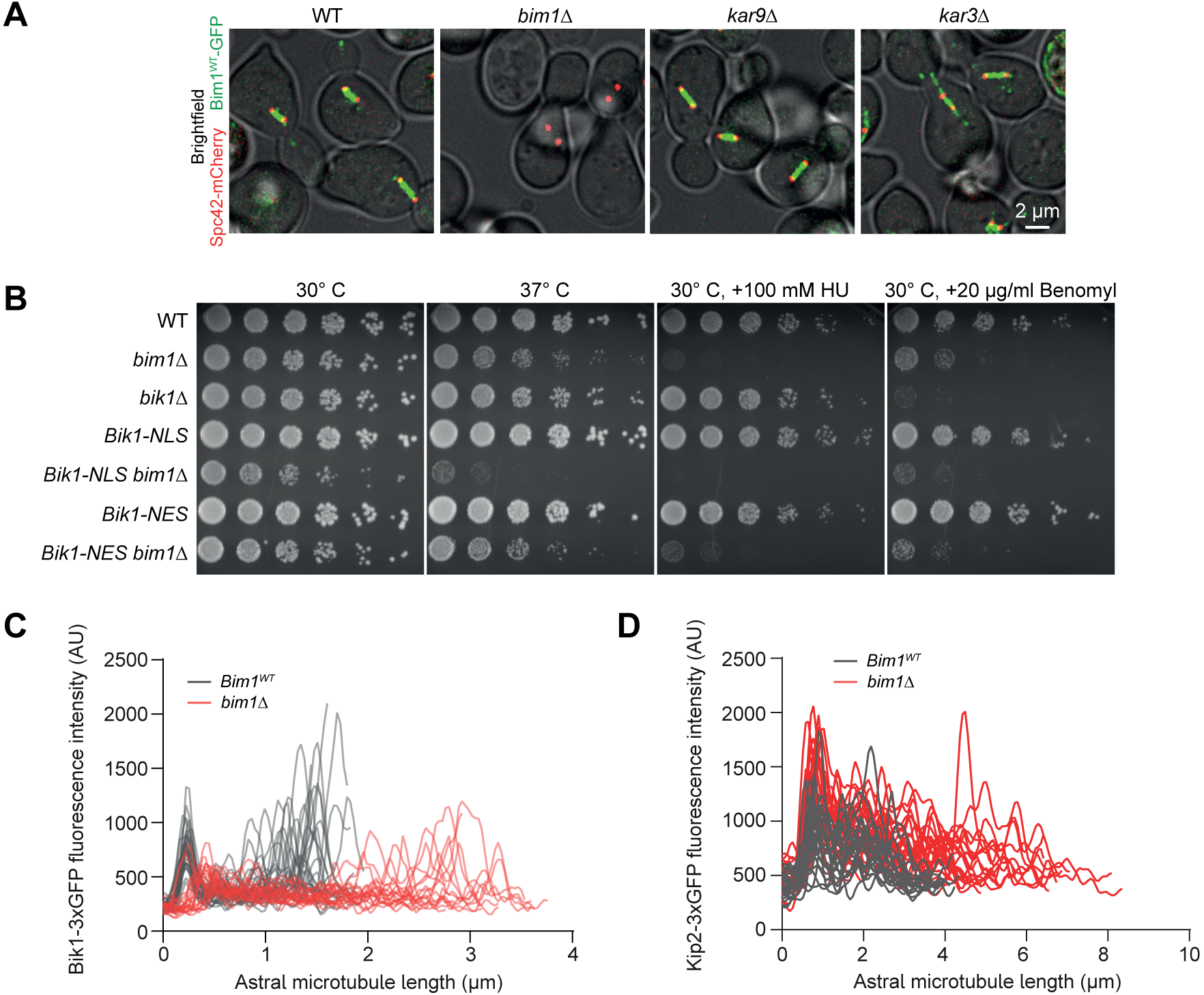
Additional analysis of *kar9Δ* and *kar3Δ* mutants. **(A)** Fluorescence microscopy analysis of localization of Bim1^WT^-GFP in deletion mutants of its major cargoes Kar9 and Kar3. Overlay of green channel (Bim1^WT^-GFP) red channel (Spc42-mCherry) with bright field image. Scale bar, 2 μm. Note the lack of cytoplasmic Bim1 signal in the *kar9Δ* mutant. **(B)** Serial dilution assay testing different BIK1 alleles alone or in combination with a *bim1Δ*. Plates were photographed after 2.5 days at 30°C, 37°C, or at 30°C in the presence of 100 mM Hydroxyurea (HU) or 20 ug/ml Benomyl. Note that *BIK1-NLS*, but not *BIK1-NES*, strongly aggravates the growth defect of the *bim1Δ* mutant. **(C)** Overlay of Bik1-3xGFP intensity on astral microtubules from 25 wild-type (grey curves) or *bim1Δ* cells (red curves). **(D)** Overlay of Kip2-3xGFP intensity on astral microtubules from 25 wild-type (grey curves) or *bim1Δ* cells (red curves).

**Supplementary Figure S6:**
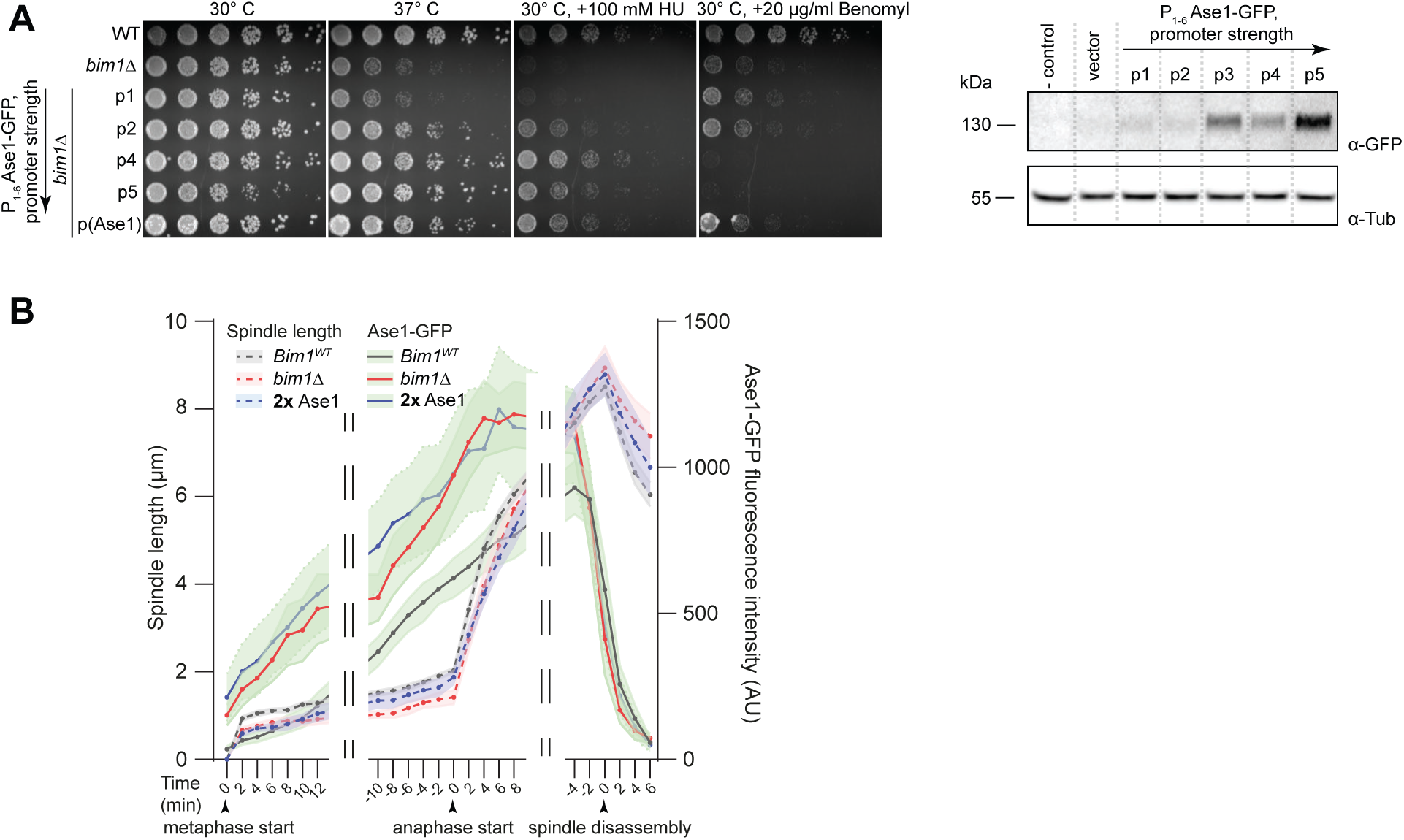
Additional characterization of the effects of increased Ase1 dosage. **(A)** Serial dilution spot assay testing growth at 30°C, 37°C or at 30°C in the presence of 100 mM Hydroxyurea or 20 μg/ml Benomyl. Plates were photographed after 2.5 days. Note the dose-dependent rescue of temperature-, hydroxyurea-sensitivity of a *bim1Δ* mutant by expression of the exogenous copy of Ase1. On the right side: western blot analysis of Ase1-GFP expression from promoters of increasing strength. α-Tubulin blot is shown as loading control. **(B)** Quantification of fluorescence intensity of Ase1-GFP (solid lines) and spindle length (dotted lines) in **2x** Ase1 *bim1Δ* strain. Data for wild-type and *bim1Δ* backgrounds is the same as in Figure 2C. Breaks in the graph are used to show correction for differences in metaphase and anaphase durations. 15 cells were analyzed for spindle size and Ase1-GFP intensity. Note that **2x** Ase1 rescues late but not early metaphase spindle length of *bim1Δ* mutant.

**Supplementary Figure S7:**
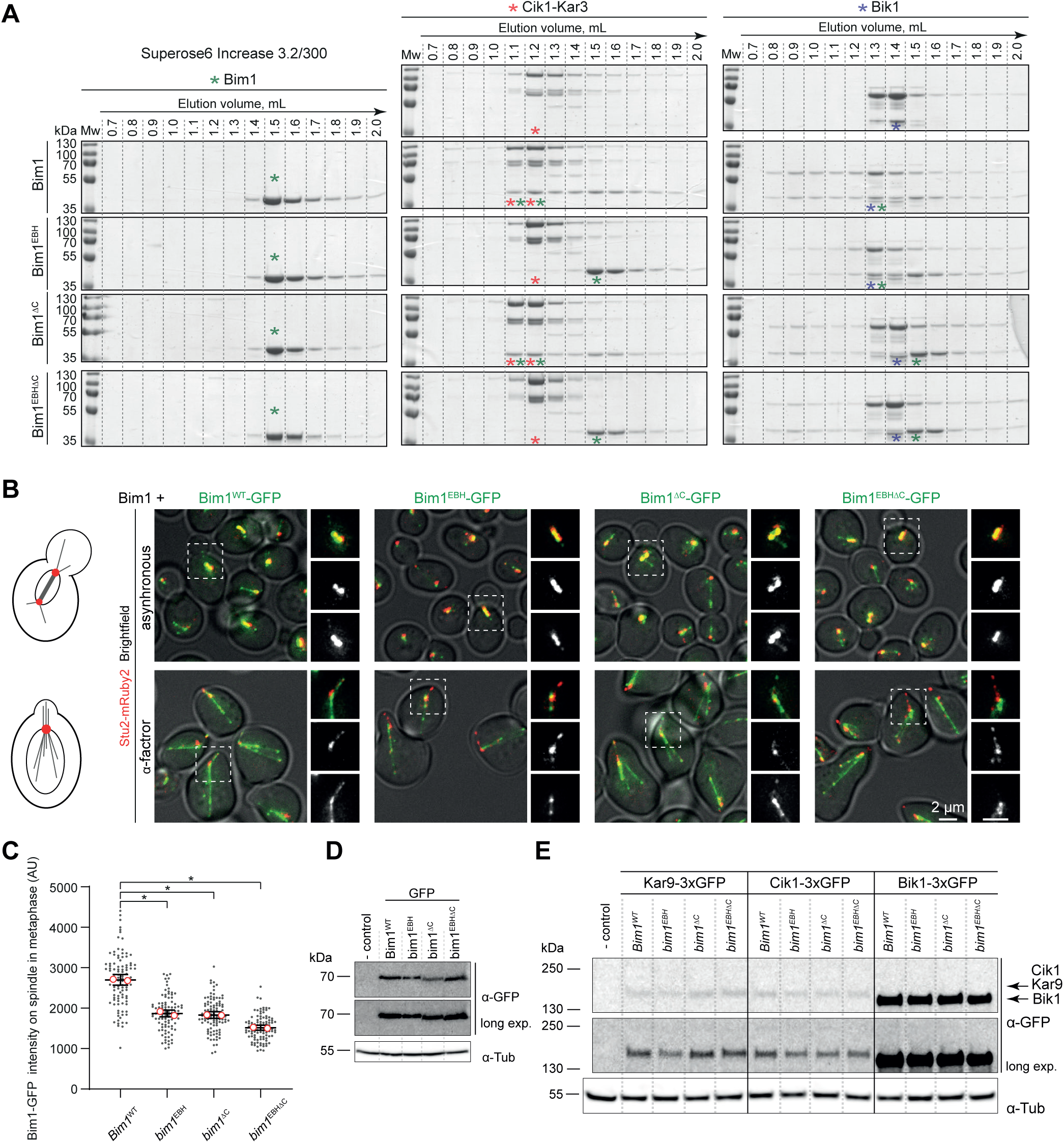
Additional characterization of *bim1^EBH^*, *bim1^ΔC^* and ***bim1^EBHΔC^* mutants *in vitro* and *in vivo***. **(A)** Size exclusion chromatography analysis of the indicated proteins and protein complexes. Sample fractions from the Superose6 Increase column were resolved on SDS-PAGE and visualized by Coomassie staining. Input concentrations of proteins were 5 µM. Positions of protein elution peaks are labelled with stars. **(B)** Live cell microscopy of Bim1 wild-type or mutant -GFP constructs in asynchronous (upper row) or α-factor arrested yeast cells. Cells also contained an unmodified *BIM1* allele. The strains additionally express Stu2-mRuby2. Left side: Overlay of fluorescence microscopy with bright field image. On the right side of every image magnification of the boxed area is shown. From top to bottom: merged image, red channel, green channel Scale bar, 2 μm. **(C)** Quantification of Bim1-GFP intensity on the metaphase spindle in the indicated strains. 100 of cells were analyzed for every condition. Small dots correspond to individual cells, open red circles represent mean values from two repeats. Error bars are mean values with 95% CIs. Asterisks denote *, P<0.0001; Welch and Brown-Forsythe ANOVA with Games and Howell post-test was used. **(D)** Western blot analysis of the expression levels of wild-type and mutant Bim1-GFP proteins. α-Tubulin was blotted as a loading control. **(E)** Protein levels of the main cargo proteins Kar9, Cik1 and Bik1 analyzed by Western blot in different *bim1* mutants as indicated. α-Tubulin blot is shown as loading control.

