## Supplementary figures and images for "Systematic analysis of microtubule plus-end networks defines EB-cargo complexes critical for mitosis in budding yeast"

### Source data F1

#### SourceData for Figure 1

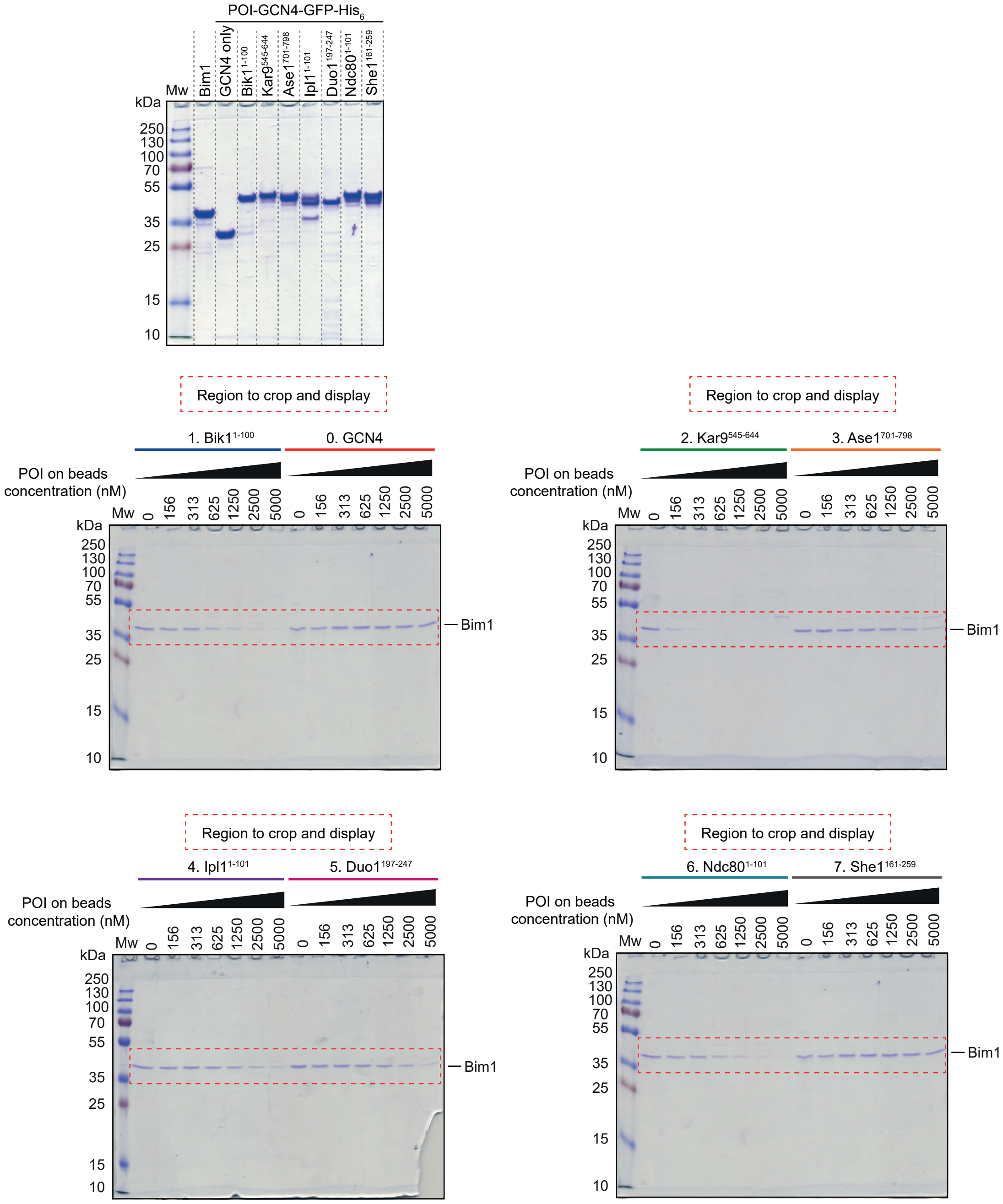

### Source data FS1

#### SourceData for Supplementary Figure 1

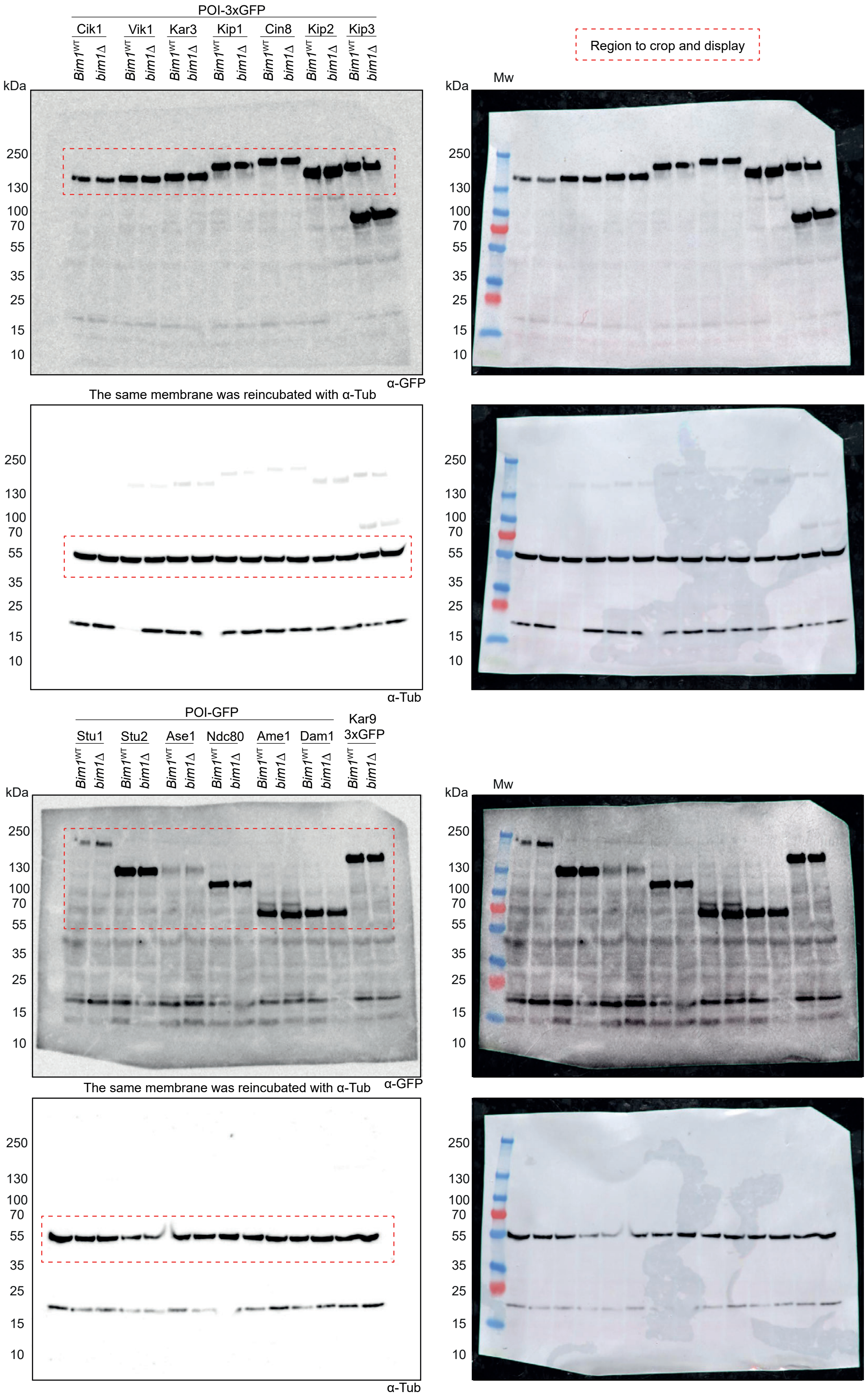
